# Shared GABA transmission pathology in dopamine agonist- and antagonist-induced dyskinesia

**DOI:** 10.1101/2023.07.26.550763

**Authors:** Yoshifumi Abe, Sho Yagishita, Hiromi Sano, Yuki Sugiura, Masanori Dantsuji, Toru Suzuki, Ayako Mochizuki, Daisuke Yoshimaru, Junichi Hata, Mami Matsumoto, Shu Taira, Hiroyoshi Takeuchi, Hideyuki Okano, Nobuhiko Ohno, Makoto Suematsu, Tomio Inoue, Atsushi Nambu, Masahiko Watanabe, Kenji F Tanaka

**Affiliations:** Division of Brain Sciences, Institute for Advanced Medical Research, Keio University School of Medicine, Tokyo, 160-8582, Japan; Laboratory of Structural Physiology, Center for Disease Biology and Integrative Medicine, Faculty of Medicine, The University of Tokyo, Tokyo, 113-0033, Japan; Division of System Neurophysiology, National Institute for Physiological Sciences, 38 Nishigonaka, Myodaiji, Okazaki, Aichi, 444-8585, Japan; Department of Physiological Sciences, SOKENDAI (The Graduate University for Advanced Studies, 38 Nishigonaka, Myodaiji, Okazaki, Aichi, 444-8585, Japan; Division of Behavioral Pharmacology, International Center for Brain Science, Fujita Health University, 1-98 Dengakugakubo, Kutsukake-cho, Toyoake, Aichi, 470-1192, Japan; Department of Biochemistry, Keio University School of Medicine, Tokyo, 160-8582, Japan; Department of Oral Physiology, Showa University School of Dentistry, 1-5-8 Hatanodai, Shinagawa-ku, Tokyo 142-8555, Japan; Division of Regenerative Medicine, The Jikei University School of Medicine, 3-25-8 Nishi-Shimbashi, Minato-ku, Tokyo 105-8461, Japan; RIKEN Center For Brain Science, 2-1 Hirosawa, Wako, Saitama, 351-0198, Japan; Graduate School of Human Health Sciences, Tokyo Metropolitan University, 7-2-10 Higashiogu, Arakawa-ku, Tokyo, 116-8551, Japan; Section of Electron Microscopy, Supportive Center for Brain Research, National Institute for Physiological Sciences, Okazaki, Aichi, 444-8585, Japan; Department of Developmental and Regenerative Neurobiology, Institute of Brain Science, Nagoya City University Graduate School of Medical Sciences, Nagoya, Aichi, 467-8601, Japan; Faculty of Food and Agricultural Sciences, Fukushima University, Kanayagawa, Fukushima 960-1248, Japan; Department of psychiatry, Keio University School of Medicine, Tokyo, 160-8582, Japan; Department of Physiology, Keio University School of Medicine, Tokyo, 160-8582, Japan; Department of Anatomy, Division of Histology and Cell Biology, Jichi Medical University School of Medicine, Shimotsuke, Tochigi, 329-0498, Japan; Division of Ultrastructural Research, National Institute for Physiological Sciences, Okazaki 444L8787, Japan; Department of Anatomy and Embryology, University of Hokkaido, Sapporo, Hokkaido, 060-8638, Japan

**Author notes:** Equal contribution (Sho Yagishita, Hiromi Sano, Yuki Sugiura, Masanori Dantsuji). Corresponding author Kenji F Tanaka 35 Shinanomachi, Shinjuku, Tokyo 160-8582, Japan.

**Keywords:** L-DOPA-induced dyskinesia, tardive dyskinesia, brain volume, structural plasticity, GPe, SNr, VGAT, GABA, medium spiny neuron, dopamine fluctuation

## Abstract

Dyskinesia is involuntary movement caused by long-term medication with dopamine-related agents: the dopamine agonist, L-DOPA, to treat Parkinson’s disease (L-DOPA-induced dyskinesia [LID]) or dopamine antagonists to treat schizophrenia (tardive dyskinesia [TD]). However, it remains unknown why distinct types of medications for distinct neuropsychiatric disorders induce similar involuntary movements. Here, we searched for a shared structural footprint using magnetic resonance imaging-based macroscopic screening and super-resolution microscopy-based microscopic identification. We identified the enlarged axon terminals of striatal medium spiny neurons in both LID and TD model mice. The striatal overexpression of vesicular gamma-aminobutyric acid transporter (VGAT) was necessary and sufficient for modeling these structural changes; VGAT levels gated the functional and behavioral alterations in dyskinesia models. Our findings indicate that lowered type 2 dopamine receptor signaling with repetitive dopamine fluctuations is a common cause of VGAT overexpression and late-onset dyskinesia formation, and that reducing dopamine fluctuation rescues dyskinesia pathology via VGAT downregulation.

**Highlights:**

- Enhancement of GABAergic transmission is a shared mechanism between LID and TD.
- VGAT levels in MSNs govern the structure and function of MSN presynaptic terminals.
- Gain and loss of VGAT function in MSNs exacerbates and ameliorates dyskinesia.
- Lowered D2 signaling with repetitive DA fluctuations causes VGAT overexpression.

**Graphical abstract:** 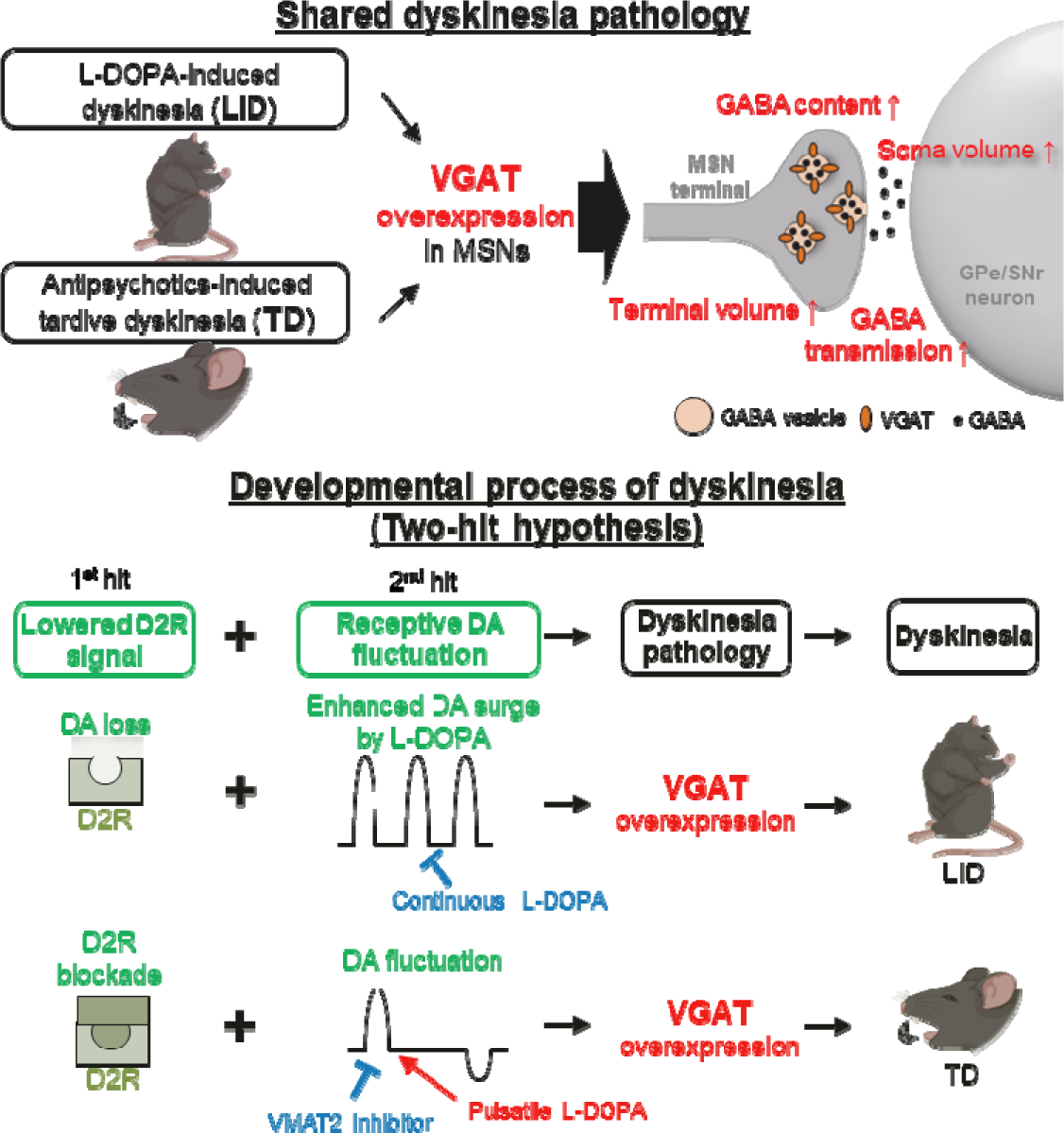

## Introduction

Dopamine-modulating medications often affect the motor system. Psychostimulants (e.g., methamphetamine and cocaine) enhance dopamine release and stimulate both type 1 and 2 dopamine receptors (D1 and D2), thus inducing hyperkinetic symptoms ^1–3^. Conventional antipsychotics (e.g., haloperidol) blockade D2 receptors and induce akinesia and rigidity, which are regarded as hypokinetic symptoms ^4, 5^. Opposite acute pharmacological effects occur for dopamine receptor agonism versus antagonism. Besides these acute effects, long-term use of either D2 agonists or antagonists can induce late-onset erratic movements known as dyskinesia.

Dyskinesia refers to involuntary movements of the face, arms, legs, or trunk and is a major side effect during drug treatments ^6, 7^. Most dyskinesias are associated with therapeutic drug treatment; there are two major types of drug-induced dyskinesias. The dopamine agonist 3,4-dihydroxy-L-phenylalanine (L-DOPA) is the first-line therapy for Parkinson’s disease (PD), and L-DOPA-induced dyskinesia (LID) often appears in PD patients after a few years of successful L-DOPA treatment ^8^. The other type of dyskinesia is tardive dyskinesia (TD), which is induced by D2-blocking agents (mainly antipsychotics) ^9, 10^. The long-term use of antipsychotics (both first and second generation) over several months can induce late-onset TD ^11^. Although both LID and TD are caused by long-term medication, it remains unknown why distinct types of medications for distinct neuropsychiatric disorders can induce similar involuntary movements. Regarding this question, we hypothesized that the development of both kinds of dyskinesia might share a common mechanism.

The genesis of dyskinesia can be understood as an irreversible brain shift from a no-dyskinesia state to a dyskinesia state after drug treatment. Such brain state changes likely involve unidentified cellular and circuitry plastic changes as well as structural changes; theoretical functional plasticity should be accompanied by identifiable structural plasticity. Pioneering anatomical studies have demonstrated structural changes in a rat model of LID; terminals of striatal medium spiny neurons (MSNs) are enlarged, thus increasing the volumes of the internal segment of the globus pallidus (GPi) and the substantia nigra pars reticulata (SNr) ^12^. This finding provide us with clues to the shared pathology of LID and TD in line with functional and structural plastic changes in the brain.

## Results

### Enlargement of inhibitory presynaptic structures in the external segment of the globus pallidus (GPe) and SNr is a shared pathological change in dyskinesias

To better understand structural plastic changes in dyskinesias, we developed a comprehensive panel of brain anatomical investigations (Figure 1A and Table S1). The panel consisted of brain-wide structural magnetic resonance imaging (MRI) screening of brain regions as well as light microscopy-, super-resolution microscopy (SRM)-, and electron microscopy (EM)-assisted identification of cellular/subcellular volume changes. We applied the panel to a well-established mouse model of LID in which hemi-parkinsonism is induced by 6-hydroxydopamine hydrobromide (6-OHDA)-mediated dopaminergic neuronal ablation and mice are then treated with L-DOPA daily for 2 weeks (Figure 1B). To validate the successful LID model ^7, 13^, we observed abnormal involuntary movements on the final day of L-DOPA administration. All 6-OHDA- and L-DOPA-treated mice had increased numbers of contralateral rotations and contralateral dystonic postures, thus confirming a successful LID model (Figure S1A, B). We also confirmed the ipsilateral ablation of striatal dopaminergic terminals using dopamine transporter (DAT) staining *post hoc* (Figure S1C). With this LID model, brain volume changes were compared using region of interest (ROI)-based volume comparisons between the ipsilateral and contralateral hemispheres of 6-OHDA injection (Figure 1D). The ipsilateral volume increased in 15 loci and decreased in nine loci (Table S2). On the basis of previous studies ^12, 14, 15^, we selected the basal ganglia for further anatomical analyses and revealed significantly increased brain volumes in the GPe, GPi, and SNr, where striatal MSNs terminate.

**Figure 1.**
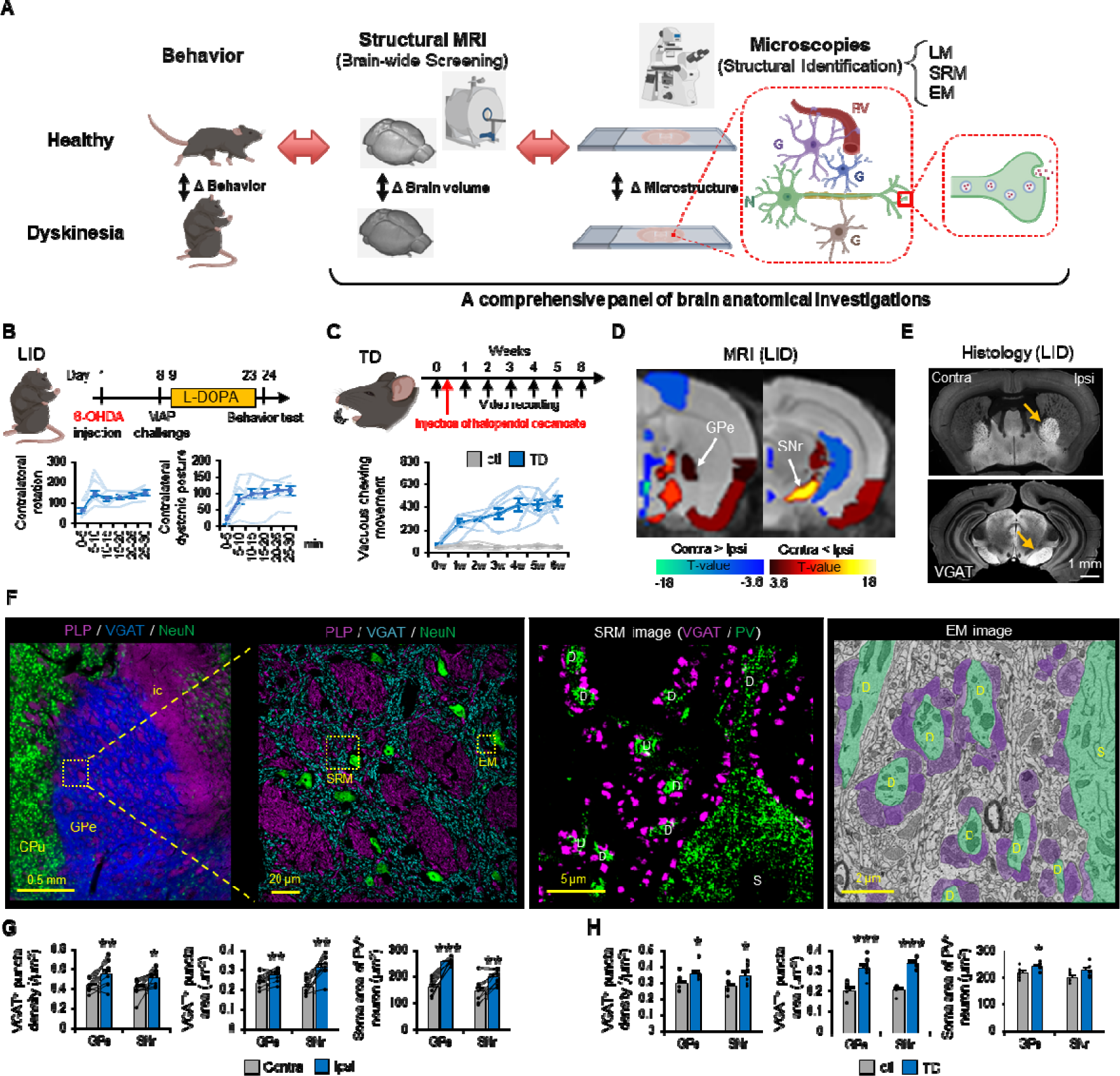
Enlargement of inhibitory presynaptic structure in the GPe and SNr is a shared pathological change in dyskinesias. (A) Schematic diagram of our research strategy. N; neuron, G; glia, BV; blood vessel. (B) Time course of the LID model mouse generation. Numbers of contralateral rotations and contralateral dystonic postures were counted every 5 minutes in LID model mice (n=6). (C) Time course of TD model mouse generation. The number of VCMs per 20 minutes was plotted every week in TD (n=6) and control (n=6) mice. (D) ROI-based brain volume changes were compared between the ipsilateral (Ipsi) and contralateral (Contra) hemispheres of LID model mice (n=8). Colored brain ROIs, showing significant changes in brain volume, were plotted (false discovery rate [FDR] corrected p<0.05). (E) VGAT immunostaining in the GPe and SNr of LID mice. The arrows show increases in Ipsi brain volumes. (F) Low- and high-magnification images of myelin proteolipid protein (PLP)/VGAT/NeuN in the GPe of control mice. ic; internal capsule. SRM images of VGAT/PV staining and an EM image of the GPe of control mice. Green indicates the soma (S) and dendrites (D) of GPe principal neurons; purple indicates MSN terminals. (G, H) VGAT^+^ density and area and soma areas of PV^+^ neurons were compared between the Contra and Ipsi hemispheres of LID mice (n=8) (G and between TD (n=5) and control (n=5) mice (H). *p<0.05. **p<0.01, ***p<0.001 (Student’s or paired t-test, p-values corrected by Bonferroni correction). Values are plotted as the mean ± standard error of the mean (SEM).

A previous microscopic study reported the enlargement of striatonigral MSN terminals in the GPi and SNr of LID model rats ^12^, thus supporting macroscopic GPi and SNr volume increases. Our MRI data also indicated engagement of the striatopallidal pathway with the GPe volume increase. To clarify this finding, we conducted immunohistochemistry for vesicular gamma-aminobutyric acid transporter (VGAT), which strongly labels γ-aminobutyric acid (GABA)ergic MSN terminals and thus delineated these nuclei (Figure 1E). The VGAT^+^ areas of the ipsilateral GPe/SNr were significantly larger than their contralateral counterparts (Figure S1D), indicating that both the striatopallidal and striatonigral pathways are involved in the volume increases of the GPe/SNr.

We then explored what types of cells and subcellular compartments were responsible for the GPe/SNr volume increases. We used a comprehensive histological panel to assess the number and size of constituent cells in the GPe/SNr and subcellular neuronal structures in both gray and white matter (Figure 1G, H; Supplementary text; Figure S1–S3; and Tables S2, S3). We found that inhibitory presynaptic structures (VGAT^+^ MSN terminals), postsynaptic structures (dendrites and soma of GPe/SNr principal neurons), and cortical myelinated axons were enlarged in the GPe/SNr of LID mice; it is possible that these anatomical changes are associated with dyskinesia development.

We next evaluated whether TD, which has a distinct etiology from LID but exhibits similar involuntary movements, shared the structural changes observed in LID. We generated a haloperidol (D2 receptor antagonist)-induced TD model by intramuscular administration of the long-acting injectable haloperidol decanoate (Figure 1C) ^16^. Typical involuntary movements of the rodent TD model are known as vacuous chewing movements (VCMs) ^16, 17^. We quantified these using two methods: the visual identification of VCMs from video data and the measurement of orofacial muscle activity by electromyogram (EMG) (Figure S1E, F). Both methods captured a gradual increase in VCMs after long-acting injectable haloperidol treatment (Figure 1C). We then examined microscopic structural changes in the corresponding orofacial region of the GPe/SNr ^18^ of the TD model mice. Inhibitory presynaptic structures in the GPe/SNr and inhibitory postsynaptic structures in the GPe were enlarged in TD mice compared with those in controls (Figure 1H and Figure S2L). However, white matter integrity of TD differed from that of LID; the diameters of penetrating myelinated axons were significantly decreased in the GPe/SNr and myelin thickness was decreased in the GPe (Figure S3C). Together, these inhibitory presynaptic and postsynaptic structural changes suggest a shared pathology between LID and TD and correlate with the development of dyskinesia.

### Increased GABA content in GPe/SNr of dyskinesia models

Previous studies have consistently reported increased mRNA expression of the 65- and 67-kDa isoforms of glutamate decarboxylase (GAD65 and GAD67, respectively; both are GABA-synthesizing enzymes) in the dopamine neuron-depleted ipsilateral striatum of LID models ^19, 20^. In addition, an imaging mass spectrometry (IMS) study revealed increased GABA content in the ipsilateral striatum and ventral pallidum of LID model mice ^21^. Because MSNs in the ventral striatum terminate in the ventral pallidum, ipsilateral MSNs in the dorsal striatum (or Caudate-Putamen; CPu) that terminate in the GPe/SNr may also contain high GABA levels in LID. To evaluate this concept, we conducted GABA IMS in the LID mouse model. As expected, GABA content was significantly increased in the ipsilateral CPu and GPe/SNr compared with that in their contralateral counterparts (Figure 2A). Similarly, the expression of GABA-related genes, including *Slc32a1* (encoding VGAT), *Gad1* (GAD67), and *Gad2* (GAD65), was significantly increased in the ipsilateral striatum (Figure S4A, B). Together, these observations suggest that increased MSN terminal volume is associated with the striatal overexpression of GABA-related genes and increased GABA content in the GPe/SNr in LID.

**Figure 2.**
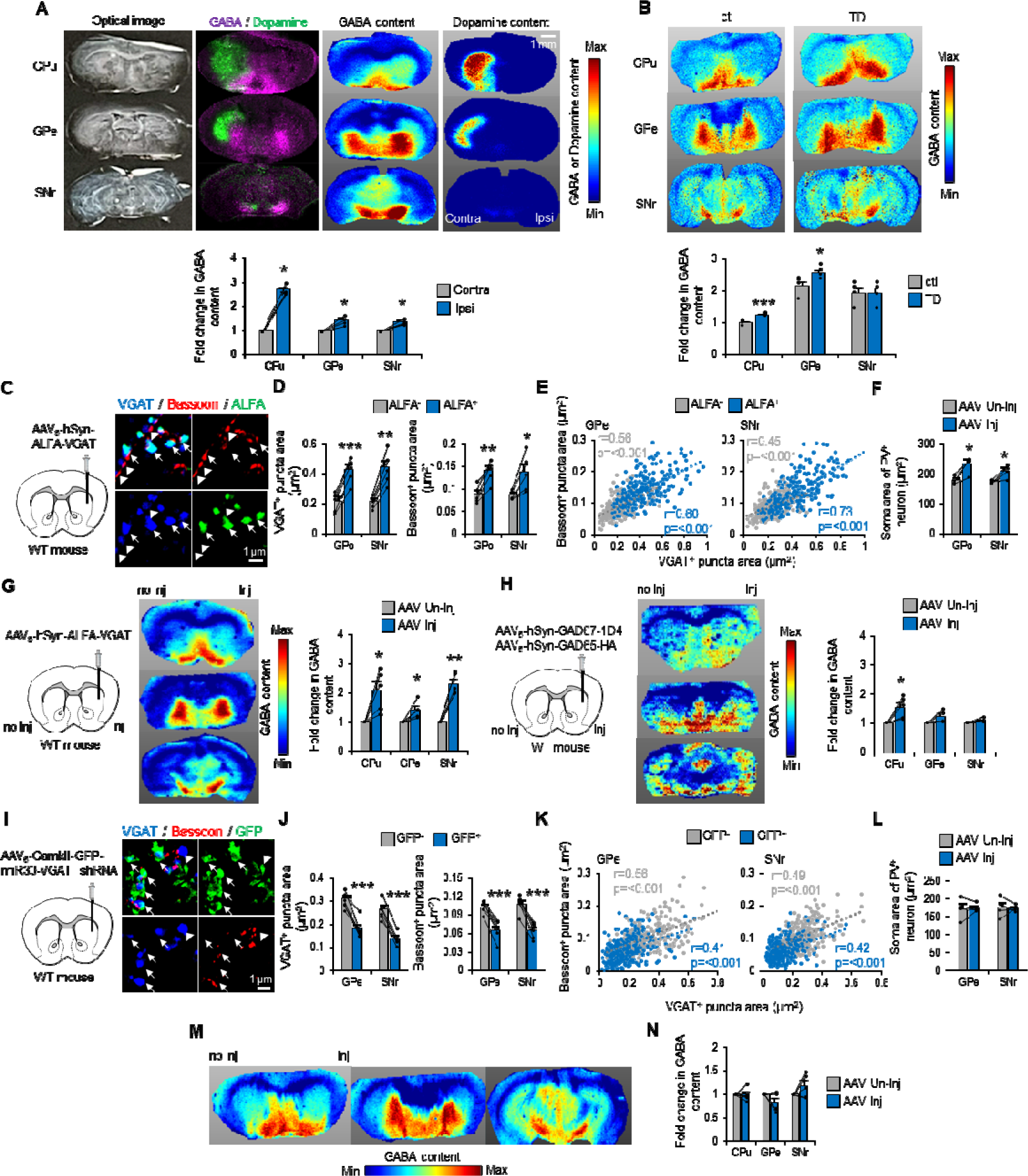
Striatal VGAT expression levels determine axon terminal size and GABA content. (A) IMS showing the optical image, GABA, dopamine, and their overlay in LID mice. Fold changes in GABA content relative to the Contra hemisphere are plotted (n=5). (B) MSI of GABA content in TD mice. Fold changes in GABA content (based on the average GABA content of each region in control mice) were plotted in control (n=4) and TD (n=4) mice. (C) SRM images of VGAT/Bassoon/ALFA in the GPe of VGAT overexpression mice. (D) Areas of VGAT^+^ and Bassoon^+^ puncta in MSN terminals were compared between ALFA^+^ and ALFA^−^ puncta in the GPe and SNr (n=5). (E) Areas of VGAT^+^/Bassoon^+^ puncta were plotted in VGAT overexpression mice. (F) Soma areas of PV^+^ neurons in the GPe and SNr were compared between the AAV injection (AAV Inj) and non-injection (AAV Un-inj) hemispheres (n=5). (G–H) IMS of GABA content in VGAT (G) and GAD67/GAD65 (H) overexpression mice. Fold changes in GABA content (relative to AAV Un-inj) plotted for AAV Inj in VGAT (n=5) and GAD67/GAD65 (n=5) overexpression mice. (I) SRM images of VGAT/Bassoon/GFP in the GPe of VGAT inhibition mice. (J) Areas of VGAT^+^ and Bassoon^+^ puncta in MSN terminals were compared between GFP^+^ and GFP^−^ puncta in the GPe and SNr (n=5). (K) Areas of VGAT^+^/Bassoon^+^ puncta were plotted in VGAT inhibition mice. (L) Soma areas of PV^+^ neurons in the GPe and SNr were compared between the AAV Inj and AAV Un-inj hemispheres (n=5). (M, N) IMS of GABA content in VGAT inhibition mice. Fold changes in GABA content (relative to AAV Un-inj) were plotted in VGAT inhibition mice (n=5). *p<0.05, **p<0.01, ***p<0.001 (Student’s or paired t-test, p-values corrected by Bonferroni’s method). Each value and the mean ± SEM are plotted.

Because enlarged MSN terminals are a common structural change in dyskinesias, we next investigated whether increased MSN terminal volume coincided with increased GABA-related gene expression in the striatum and increased GABA content in the GPe/SNr in TD. Although *Gad2* mRNA levels were significantly decreased in TD mice compared with those in controls, *Gad1* and *Slc32a1* mRNA levels were comparable (Figure S4B). Moreover, IMS revealed increased GABA content in the GPe of TD mice (Figure 2B). These data suggest that local VGAT protein levels govern the size and GABA content of MSN terminals but not the mRNA expression levels of GABA-synthesizing enzymes.

### Striatal VGAT overexpression causes the increase in MSN termina size and GABA content

Molecular and biochemical findings from the TD mice suggested that the packaging of GABA into presynaptic vesicles, rather than its synthesis, is key to structural changes in dyskinesias. To address whether VGAT overexpression *per se* recapitulated a shared pathology between LID and TD, we compared the effect of VGAT overexpression alone (mimicking TD) with that of VGAT/GAD67/GAD65 overexpression (mimicking LID) on MSN terminal size. We generated adeno-associated virus (AAV) vectors expressing VGAT, GAD67, or GAD65. A single AAV or mixture of all three AAV vectors was injected into the right dorsal striatum of wild-type (WT) mice. After AAV-mediated ALFA-tagged VGAT overexpression in MSNs, ALFA^+^/VGAT^+^ puncta were significantly enlarged in the GPe and SNr compared with ALFA^−^/VGAT^+^ puncta (Figure 2C, D). Furthermore, the presynaptic active zone (ALFA^+^/Bassoon^+^ puncta) was significantly enlarged compared with ALFA^−^/Bassoon^+^ puncta (Figure 2D, E). Striatal VGAT overexpression induced increases in the soma area of parvalbumin (PV)^+^ principal neurons (Figure 2F) as well as in the axon diameter and myelin thickness of cortical myelinated axons in the GPe (Figure S4C). With triple GABA-related gene overexpression, similar structural changes were observed (Figure S4E-G), indicating that striatal VGAT overexpression is sufficient to induce dyskinesia-relevant MSN terminal volume increases.

We then evaluated whether striatal VGAT overexpression *per se* (rather than striatal GAD overexpression) increased GABA content in the GPe/SNr. GABA content was significantly increased in the GPe, SNr, and CPu after AAV-mediated striatal VGAT overexpression compared with that in the uninjected hemisphere (Figure 2G). In contrast, GABA content was not upregulated in the GPe/SNr after striatal GAD65/67 overexpression, although GABA content was upregulated in the CPu (Figure 2H). These data indicate that the packaging of GABA into presynaptic vesicles, rather than its synthesis, increases GABA content in MSN terminals.

We next conducted a striatal loss-of-function study using short hairpin RNA (shRNA) targeting *Slc32a1* (VGAT) mRNA. We generated AAV carrying enhanced green fluorescent protein (GFP)-miR30-VGAT shRNA ^22^ and investigated whether shRNA-mediated striatal VGAT inhibition decreased MSN terminal size in WT mice. As expected, GFP^+^/VGAT^+^ and GFP^+^/Bassoon^+^ puncta size in the GPe/SNr were decreased by VGAT loss of function compared with GFP^−^/VGAT^+^ or GFP^−^/Bassoon^+^ puncta size (Figure 2I–K). Together with gain-of-function data, this suggests that VGAT levels determine presynaptic structure size in MSNs. In contrast, striatal VGAT loss of function did not induce structural changes in GPe/SNr principal neurons (Figure 2L) or penetrating myelinated axons (Figure. S4D) and did not reduce GABA content (Figure 2M, N) in WT mice.

### Striatal VGAT overexpression enhance GABA transmission from MSNs

Given that we have previously demonstrated enhanced GABA transmission from MSNs in the GPe and SNr in LID ^23^, we examined whether GABA transmission was also enhanced in TD and whether VGAT overexpression enhanced GABA transmission. In our experimental setup, we recorded single unit activity in orofacial and forelimb regions of the GPe and SNr and examined their responses to electrical stimulation of the cerebral cortex (Cx). The typical response pattern of GPe and SNr neurons is a triphasic response composed of early excitation (i), inhibition (ii), and late excitation (iii) (Figure 3A) ^23^. Each component in the GPe is mediated by the Cx–subthalamic nucleus (STN)–GPe (i), Cx–D2 MSN (striatopallidal)–GPe (ii), and Cx–D2 MSN–GPe–STN–GPe (iii) pathways. Each component in the SNr is mediated by the Cx–STN–SNr (i), Cx–D1 MSN (striatonigral)–SNr (ii), and Cx–D2 MSN–GPe–STN–SNr (iii) pathways. Alterations in Cx-evoked triphasic responses can therefore highlight neurotransmission changes through each basal ganglia pathway.

**Figure 3.**
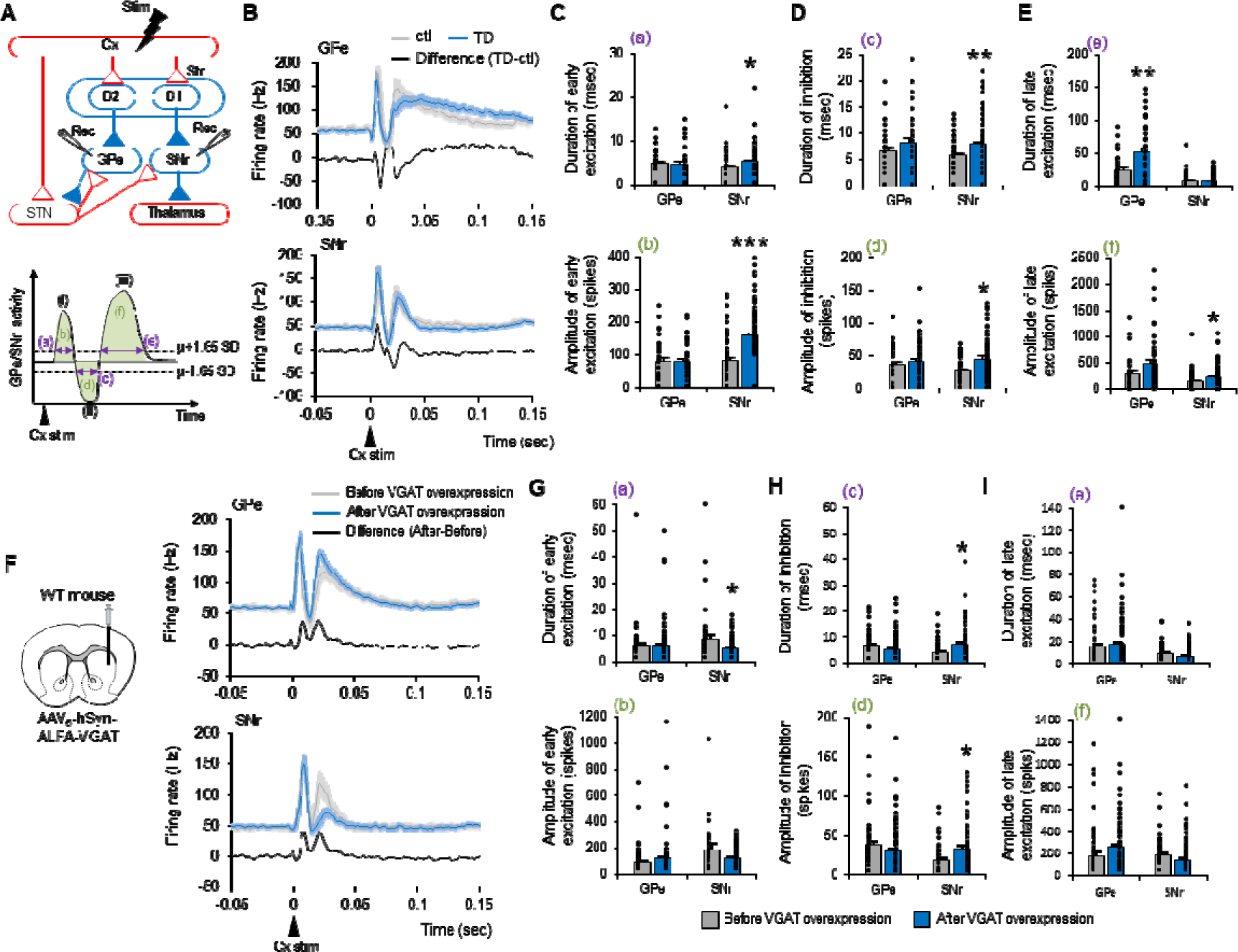
Striatal VGAT overexpression enhances GABA transmission from MSNs. (A) Stimulation (Cx) and recording (GPe or SNr) sites are depicted with basal ganglia circuitry. Red and blue lines represent glutamatergic excitatory and GABAergic inhibitory projections, respectively. In the striatum (Str), D1 and D2 represent D1-MSN and D2-MSN, respectively. Cx-evoked responses in the GPe or SNr are typically composed of early excitation (i), inhibition (ii), and late excitation (iii). Purple lines represent the durations of the three responses (a, c, e), and green areas represent the amplitudes of the three responses (b, d, f). (B) Averaged PSTHs of Cx-evoked responses in the GPe (control, n=30 neurons, gray; TD, n=38 neurons, blue; from four mice) and SNr (control, n=52 neurons; TD, n=89 neurons; from four mice). Algebraic differences between TD and controls are also indicated (black). (C–E) Durations and amplitudes of early excitation, inhibition, and late excitation responses were compared between control and TD mice. (F) Single-unit recording was performed before and after VGAT overexpression. Averaged PSTHs of Cx-evoked responses in the GPe (before n=52 neurons, after n=123 neurons, from four mice) or SNr (before n=37 neurons, after n=61 neurons, from four mice) before and after striatal VGAT overexpression in WT mice. Differences in averaged PSTHs before and after overexpression were calculated, as were algebraic differences between before and after overexpression. (G–I) Duration and amplitude of early excitation, inhibition, and late excitation responses were compared before and after VGAT overexpression. *p<0.05, **p<0.01 (Student’s t-test, p-values corrected by Bonferroni’s correction). Values of each mouse and the mean ± SEM are plotted.

Averaged peristimulus time histograms (PSTHs) of GPe and SNr neurons between control and TD mice and their algebraic differences were plotted (Figure 3B). Both the duration and amplitude of inhibition (ii) were significantly increased in the SNr of TD mice (Figure 3D), indicating increased GABA release from striatonigral MSNs. The duration of late excitation (iii) was significantly increased in the GPe and its amplitude was increased in the SNr (Figure 3E), suggesting increased GABA release from striatopallidal MSNs. Early excitation (i), which is irrelevant to MSN GABA release, was also significantly increased in the SNr (Figure 3C), possibly indicating structural changes in VGluT2^+^ excitatory terminals (Figure S3B). These results indicate increased GABA release from striatonigral and striatopallidal MSNs in the TD mice.

We next assessed the effects of striatal VGAT overexpression *per se* on GABA release from MSN terminals. Averaged PSTHs of GPe and SNr responses were compared before and after AAV injection (Figure 3F). The duration and amplitude of inhibitory responses were increased in the SNr (Figure 3H), indicating increased GABA release from striatonigral MSNs. Although the duration of late excitation was comparable between before and after AAV injection, the amplitude of late excitation tended to be increased in the GPe (p<0.1) (Figure 3I), suggesting increased GABA release from striatopallidal MSNs. These data indicate that striatal VGAT overexpression results in the structural and functional augmentation of GABA release from presynaptic MSN terminals.

### Striatal VGAT expression levels gate the severity of dyskinesia

Striatal VGAT levels determine MSN terminal size and GABA transmission, and striatal VGAT overexpression may be a common mechanism of both types of dyskinesia. To better understand this phenomenon, we conducted VGAT gain- and loss-of-function studies in the two dyskinesia models and evaluated the degree of involuntary movement. In the LID model, a VGAT- or GFP-expressing AAV vector was injected into the right side of the dorsal striatum; 6-OHDA was injected into the same side 2 weeks later (Figure 4A). VGAT overexpression did not induce rotational behaviors or dystonic postures before L-DOPA administration; however, abnormal involuntary movements were observed even on the first day of L-DOPA administration and remained significantly more common on the last day (Figure 4B). In the TD model, VGAT- or GFP-expressing AAV vectors were injected into the bilateral striatum 3 weeks before the administration of haloperidol decanoate (Figure 4C). VGAT overexpression induced vacuous chewing movements (VCMs) without haloperidol administration (Figure 4D), which was unexpected, and enhanced VCMs with haloperidol administration. In contrast, GAD65/67 overexpression did not enhance VCMs (Figure S5A). In both dyskinesia models, increased VGAT gene expression exacerbated involuntary movements.

**Figure 4.**
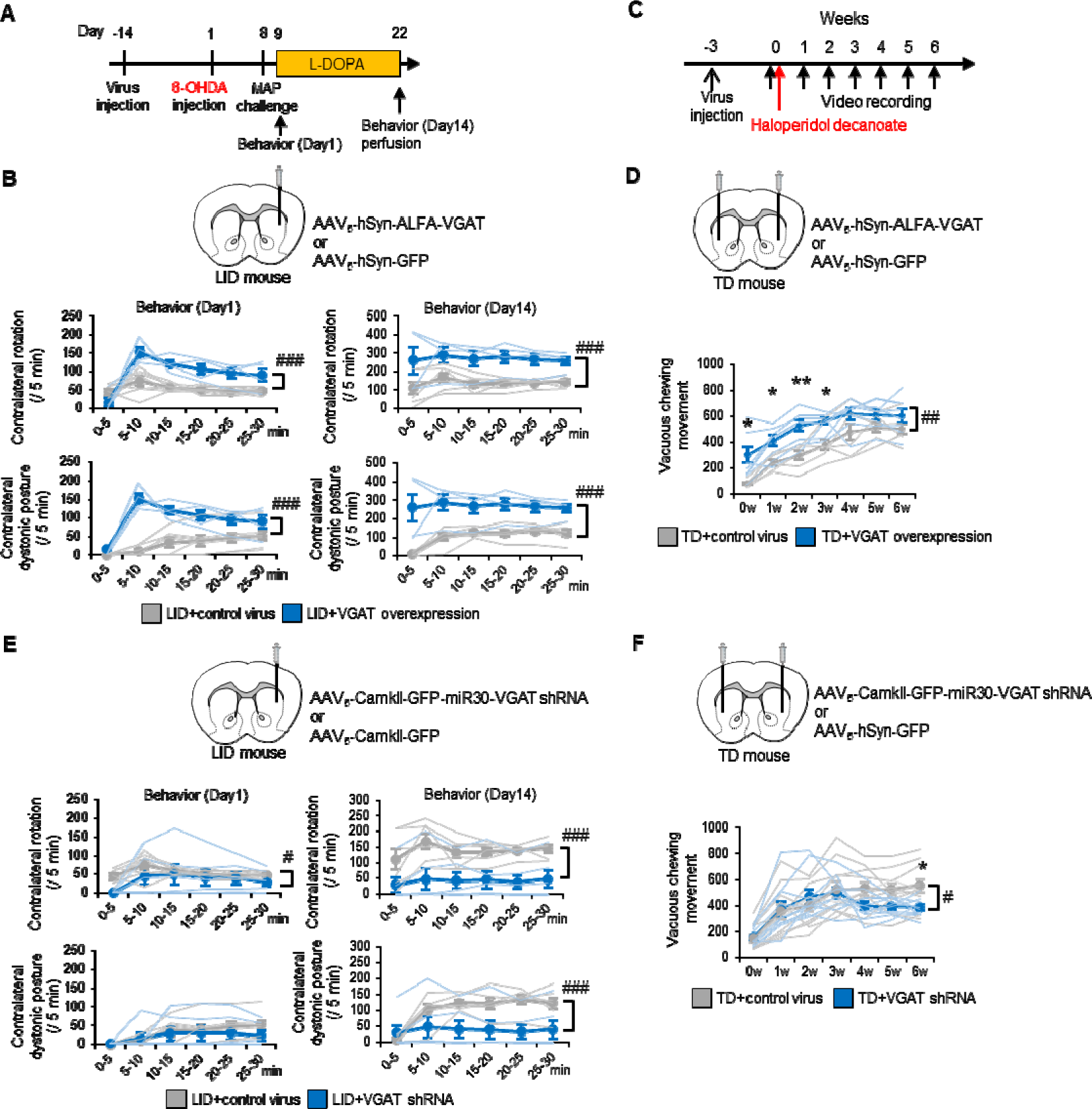
Striatal VGAT expression levels gate the severity of dyskinesia. (A) AAV vectors were injected into the right dorsal striatum (ipsilateral to the 6-OHDA injection) of LID mice 2 weeks before 6-OHDA injection; L-DOPA was then injected for 2 weeks. Mouse behavior was observed on the first and final days of L-DOPA injection. (B) Numbers of contralateral rotations and contralateral dystonic postures were compared between LID with VGAT overexpression (n=4) and LID with control AAV (n=7) mice. (C) AAV vectors were injected into the bilateral dorsal striatum of TD mice 3 weeks before haloperidol decanoate injection. (D) The number of VCMs was compared between TD with control AAV (n=6) and TD with VGAT overexpression (n=6) mice. (E) Numbers of contralateral rotations and contralateral dystonic postures were compared between LID with VGAT shRNA (n=6) and LID with control AAV (n=7) mice. (F) Numbers of VCMs were compared between TD with VGAT shRNA (n=10) and TD with control AAV (n=11) mice. #p<0.05, ##p<0.01, ###p<0.001 (two-way repeated analysis of variance [ANOVA]). *p<0.05, **p<0.01 (Student’s t-test, p-values corrected by Bonferroni’s correction). Values of each mouse and the mean ± SEM are plotted.

We next attempted to mitigate dyskinesia by targeting the overexpressed striatal VGAT. In the LID model, prior to LID induction, AAV carrying VGAT shRNA or GFP was injected into the ipsilateral dorsal striatum. During L-DOPA administration, LID behaviors were markedly decreased (Figure 4E). Notably, ipsilateral VGAT^+^ MSN terminal size was decreased rather than increased (Figure S5B, C), and the PV^+^ soma areas of GPe and SNr neurons were not enlarged (Figure S5D). These changes were consistent with the loss-of-function study in WT mice (Figure 2I–L). In the TD model, AAVs were bilaterally injected into the dorsal striatum before haloperidol treatment. Visually identified VCMs were decreased 3 weeks after the haloperidol injection (Figure 4F) and the suppression of increased orofacial electromyography (EMG) activity was identified before 3 weeks (Figure S5J). The suppression of enlarged VGAT^+^ MSN terminals and enlarged PV^+^ soma in GPe neurons was also identified (Figure S5F-I). These results indicate that striatal VGAT levels govern the pathology and pathophysiology of both TD and LID.

### Two-hit model of dyskinesia formation

We next speculated upon the events that might lead to a shared structural footprint in both LID and TD. Clinical observations suggest that continuous L-DOPA treatment, such as Duodopa intestinal infusion, reduces LID risk compared with conventional oral L-DOPA administration ^24, 25^, which may induce dopamine surges. We therefore investigated the constructive role of pulsatile dopamine fluctuation on dyskinesia formation. As previously demonstrated in the LID rodent model ^26^, we continuously administered L-DOPA after 6-OHDA treatment using a subcutaneous slow-release L-DOPA pellet ^27^ and found no dyskinesia (Figure 5A). To rule out the possibility that insufficient L-DOPA treatment failed to produce dyskinesia, we examined whether hemi-parkinsonism was treated with this system. We trained mice to perform a single-forelimb reaching task and evaluated forelimb movements ^28^. Continuous L-DOPA administration markedly improved 6-OHDA-mediated impaired reaching behavior, confirming the successful L-DOPA treatment of hemi-parkinsonism (Figure S6A, B). In this condition, there were no increases in the VGAT^+^ MSN terminal size or PV^+^ soma area of GPe and SNr principal neurons (Figure 5B, C), indicating that continuous dopamine supplementation does not cause LID-associated structural changes or abnormal involuntary movements.

**Figure 5.**
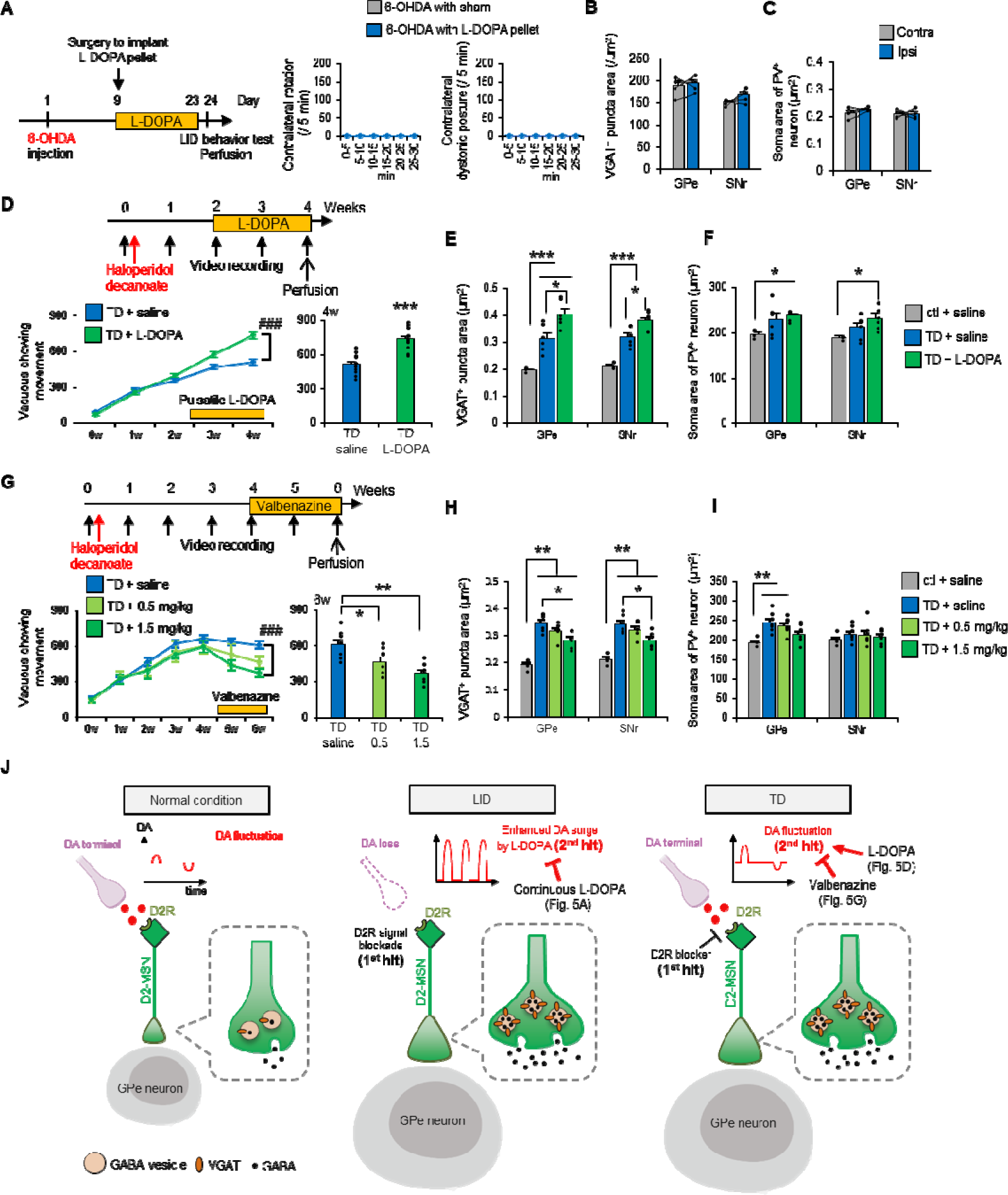
Lowered dopamine receptor type 2 signaling with repetitive dopamine fluctuations induces VGAT overexpression and dyskinesia. (A) Experimental time course of continuous L-DOPA administration. The numbers of contralateral rotations and contralateral dystonic postures were counted in L-DOPA-treated (n=4) and sham (n=3) mice. (B, C) The area of VGAT^+^ puncta of MSN terminals and soma area of PV^+^ neurons were compared between the Ipsi and Contra hemispheres of mice with continuous L-DOPA (n=4). (D) Experimental time course for the pulsatile administration of L-DOPA (daily, intraperitoneal) in TD mice. Numbers of VCMs were plotted every week and the 4-weeks period in TD mice with saline (n=12) and L-DOPA (n=12) treatment. (E, F) The area of VGAT^+^ puncta of MSN terminals and soma area of PV^+^ neurons were compared among control mice with saline (n=3), TD mice with saline (n=6), and TD mice with L-DOPA (n=6). (G) Experimental time course of daily valbenazine administration (0.5 or 1.5 mg/kg, oral administration) in TD mice. Numbers of VCMs were plotted every week and the 6-weeks period in TD mice with saline (n=8) and valbenazine (n=8 for each dose) treatment. (H, I) The area of VGAT^+^ puncta of MSN terminals and soma area of PV^+^ neurons were compared among control mice (n=4), TD mice with saline (n=7), and TD mice with valbenazine (n=7 for each dose). (J) Schematic diagram of proposed dyskinesia pathology. In LID, the ablation of dopaminergic neurons (1^st^ hit) and evoked dopamine (DA) surges by L-DOPA administration (2^nd^ hit) increases striatal VGAT expression, resulting in volume increases in MSN terminals and soma of GPe/SNr neurons, and increased GABA content and transmission. In TD, blocking of D2 receptors (D2R; 1^st^ hit) and physiological dopamine fluctuations (2^nd^ hit) induce similar pathophysiology to LID. The increased and decreased amplitudes of dopamine fluctuations induced by L-DOPA and valbenazine, respectively, led to exacerbated (with L-DOPA) and ameliorated (with valbenazine) dyskinesia pathology. *p<0.05, **p<0.01, ***p<0.001 (Student’s t-test, p-values corrected by Bonferroni correction). Values are plotted as the mean ± SEM.

Because ambient striatal dopamine concentrations are sufficient to occupy high-affinity D2 receptors but not low-affinity D1 receptors ^29, 30^, the ablation of dopamine neurons in PD leads to long-term vacant D2 receptors and a resultant continuous loss of D2 signaling. Pharmacological D2 receptor antagonism decreases D2 signaling; long-term D2 blockade, as occurs with long-term antipsychotic use, can therefore also be regarded as a continuous loss of D2 signaling. Accordingly, LID and TD likely share a common etiology in terms of D2 signaling. In the case of LID, in addition to the initial hit (a lasting loss of D2 signaling), L-DOPA-mediated dopamine fluctuation (a second hit) leads to dyskinesia. In the case of TD, physiological dopamine fluctuation (that is, a dopamine surge or dip in response to salient stimuli in daily life) may function as the second hit (Figure 5J).

To experimentally evaluate the relevance of the second hit in TD, we artificially enhanced dopamine fluctuations via pulsatile L-DOPA administration under continuous haloperidol treatment. After L-DOPA administration, the number of VCMs (Figure 5D) and the size of VGAT^+^ MSN terminals and PV^+^ soma in GPe/SNr neurons (Figure 5E, F) were significantly increased. In contrast, 2 weeks of pulsatile L-DOPA administration without a loss of D2 signaling did not increase the size of VGAT^+^ MSN terminals (Figure S2A). Together, these results support the two-hit model of TD.

We finally addressed whether and how the vesicular monoamine transporter 2 (VMAT2) inhibitor valbenazine might mitigate VCMs in the TD model ^31–33^. After VCM acquisition, mice were administered valbenazine for 2 weeks (Figure 5G). VCMs were dose-dependently reduced and VGAT^+^ puncta size was normalized (Figure 5H), suggesting that the therapeutic mechanism of valbenazine is mediated by a reduction in striatal VGAT. Because this VMAT2 inhibitor also downregulates dopamine release ^34, 35^, it may also lower the amplitude of dopamine fluctuations triggered by daily salient stimuli, thus reducing the impact of the second hit.

## Discussion

Motor learning relies on changes in brain function. This functional plasticity is likely accompanied by brain structural changes ^36, 37^. It is therefore expected that the acquisition of involuntary movements, such as in LID and TD, also relies on functional and structural plasticity. We aimed to identify structural plasticity and address the molecular mechanisms leading to structural changes with a focus on structural–functional correlations. The current study revealed the enlargement of MSN presynaptic terminals and GPe/SNr principal neurons in LID and TD. These structural changes correlated with increased GABA content and enhanced GABA transmission at MSN terminals.

The pathogenesis of dyskinesia takes a relatively long time, and it is challenging to follow its development. We believe that structural changes reflect functional changes at every moment. In turn, functional changes during the disease process can be identified by visualizing the trajectory of structural plastic changes. Our results suggest that MSN terminal size mirrors dopamine fluctuation during dyskinesia progression. MSN terminal enlargement was induced by pulsatile but not continuous administration of L-DOPA in the LID model, and by the L-DOPA administration-induced enhancement of dopamine fluctuation in the TD model. In addition, MSN terminal enlargement was reduced by the administration of valbenazine, the only currently approved drug for TD, in the TD model. Together, these findings suggest that structural changes in MSN terminals may be a marker of the dyskinesia developmental process.

Two steps are likely necessary to develop dyskinesia (the two-hit model of dyskinesia, Figure 5J). This model is easy to apply to LID because dopaminergic neuronal loss (first hit) and pulsatile L-DOPA administration (second hit) are clearly identifiable. The application of the two-hit model to TD is more challenging; however, the following two clinical observations support this model. Patients with TD receive dopamine D2 receptor-blocking agents, such as antipsychotics (first hit). Given that substance abuse/dependence is a risk factor of TD ^38^, the second hit may be dopamine fluctuation induced by substance use. The use of addictive substances such as psychostimulants (e.g., cocaine and amphetamine) evokes an increase in dopamine release, leading to a hedonic response. Psychological stress, the main cause of substance use, may also increase dopamine level fluctuations. Indeed, positron emission tomography studies have reported increased dopamine after psychological stress ^39^ and a positive association between psychological stress and dopamine release in healthy subjects ^40^. Taken together, the two-hit dyskinesia model may apply to LID, TD, and beyond.

What corresponds to structural plasticity in inhibitory synapses? In the case of excitatory post-synapses, structural plasticity corresponds to newly formed dendritic spines and the increased volume of existing spines; in both cases, the surface area at the postsynaptic neuron increases. In contrast, inhibitory synapses do not form postsynaptic dendritic spines. Thus, if the number and size of the presynaptic active zone increases, postsynaptic neurons would have to gain receptive sites paired with the active zones. We demonstrated that AAV-mediated VGAT overexpression in MSNs increased Bassoon^+^ presynaptic structures. We also found increased gephyrin^+^ postsynaptic structures in GPe/SNr neurons; expanding dendritic and somal volumes coincided with the increased postsynaptic structure. Accordingly, we hypothesize that structural plasticity in MSN GPe/SNr inhibitory synapses occurs in the following order: 1) MSN axon terminal volume increases in a VGAT-dependent manner; 2) presynaptic active zone increases; 3) postsynaptic sites are enlarged; and 4) proximal dendrites and soma, where inhibitory synapses form, are enlarged because of the storage of expanding presynaptic sites in GPe/SNr neurons. Given that the structural basis of inhibitory synaptic plasticity is largely unknown, our discovery and proposal shed light on the possible mechanisms of inhibitory synaptic plasticity.

The present study has the following clinical significance. First, we clearly showed volume increases in the GPe and SNr of LID and TD model mice. Similar changes are observed in the GPe, SNr, and probably GPi of LID and TD patients. These volume changes may therefore be a biomarker of dyskinesia. Second, the suppression of striatal VGAT activity ameliorated morphological changes and dyskinesia. Striatal VGAT may thus be a therapeutic target (e.g., striatal VGAT suppression by viral vectors or drugs).

In summary, we found a shared pathology between LID and TD: increased volumes of presynaptic MSN terminals and postsynaptic soma in GPe/SNr neurons. Striatal overexpression of VGAT was necessary and sufficient to induce these structural signatures that correlated with functional changes such as the increased GABA content and GABA transmission at MSN terminals. We propose that a long-term reduction of MSN D2 signaling with dopamine fluctuation initiates VGAT-dependent dyskinesia development.

## Acknowledgments

We thank Mrs. Atsuko Imai and Noriko Hattori (National Institute for Physiological Sciences) for technical assistance. We thank the Collaborative Research Resources, Keio University School of Medicine, for technical assistance with the Zeiss ELYRA 3D-SIM system. We also thank Bronwen Gardner, PhD, from Edanz (https://jp.edanz.com/ac) for editing a draft of this manuscript. Figure 1A and Figure. 5J were created with BioRender.com. This work was supported by a Grant-in-Aid from the Japan Society for the Promotion of Science (JSPS) under grant numbers 22K15215 (Y.A.), JP21H02594 (S.Y.), JP21H05176 (S.Y.), and 21K07257 (H.S.), Moonshot R&D from the Japan Science and Technology Agency (JST) under grant number JPMJMS2021 (S.Y.), Takeda Science Foundation (Y.A.), Astellas Foundation for Research on Metabolic Disorders (Y.A.), a grant from Brain Mapping by Integrated Neurotechnologies for Disease Studies (Brain/MINDS) by the Japan Agency for Medical Research and Development (AMED) under grant numbers JP22dm0207069 (K.F.T, S.Y.), JP22dm0207001 (H.O.), and JP21dm0207115 (A.N.), and a Grant-in-Aid for Scientific Research on Innovative Areas-Resource and technical support platforms for promoting research of Advanced Bioimaging Support under grant number JP16H06280.

## Author Contributions

Conceptualization, Y.A. and K.F.T.; Methodology, Y.A. and K.F.T.; Validation, Y.A. and K.F.T.; Writing – Original Draft, Y.A.; Writing – Review & Editing, Y.A., H.T., A.N., and K.F.T.; Visualization, Y.A. and K.F.T.; Supervision, K.F.T.; Project Administration, K.F.T.; Funding Acquisition, Y.A., S.Y., H.S., H.O., A.N., and K.F.T. Y.A. performed all experiments and conducted the histological and behavioral data analysis. S.Y. supplied all AAVs. H.S. and A.T. obtained and analyzed the single-unit recording data. Y.S., T.S., S.T., and M.S. obtained the GABA mass spectrometry imaging data. M.D., A.M., and T.I. obtained and analyzed the EMG data in TD mice. D.Y., J.H., and H.O obtained the MRI images. M.M and N.O. obtained the EM images. M.W. supplied antibodies.

## STAR Methods

### Resource availability

#### Lead contact

Further information and requests for resources should be directed to and will be fulfilled by the lead contact, Kenji F Tanaka.

#### Materials availability

This study did not generate new unique reagents.

#### Data and Code availability

This paper does not report original datasets and original code. Any additional information required to reanalyze the data reported in this paper is available from the lead contact upon request.

### Experimental model and subject details

#### Animals

All animal procedures were conducted in accordance with the National Institutes of Health Guide for the Care and Use of Laboratory Animals and approved by the Animal Research Committee of Keio University School of Medicine (A2022-029, for the behavioral study), the Institutional Animal Care and Use Committees of the National Institutes of Natural Sciences (for the single-unit recording of neuronal activity), or Showa University (14018, for the EMG recording in TD mice). Experiments were performed using 8- to 12-week-old male and female mice. All mice were maintained on a 12-h/12-h light/dark cycle (lights on at 08:00) and behavioral experiments were conducted during the light phase. Female ICR mice (7–8 weeks old) and male and female C57BL/6j mice (7–8 weeks old) were purchased from Oriental Yeast Co., Ltd. (Tokyo, Japan).

## Method details

### Generating LID model mice

Desipramine hydrochloride (25 mg/kg, intraperitoneally [i.p.]) was administered to mice 30 minutes before 6-OHDA or vehicle infusion. Mice were anesthetized with a mixture of ketamine and xylazine (i.p., 100 mg/kg and 10 mg/kg, respectively) and fixed in a stereotaxic apparatus (SM-15, Narishige Scientific Instrument, Tokyo, Japan). The skull surface was exposed and the periosteum and blood were removed. Next, 1 μL of 5 μg/μL 6-OHDA or vehicle was unilaterally injected into the right medial forebrain bundle through a small hole (−1.2 mm anteroposterior and 1.2 mm mediolateral from bregma and −4.7 mm dorsoventral from the brain surface, according to the atlas of Paxinos and Franklin, 2008) using a glass micropipette connected to a Nanoliter 2020 injector (World Precision Instruments, Sarasota, FL, USA) to lesion the nigrostriatal dopaminergic neurons. 6-OHDA was dissolved in saline containing 0.02% ascorbic acid. The mice were allowed to recover for 1 week before undergoing a methamphetamine challenge to select mice in whom more than 90% of the dopaminergic terminals were successfully lesioned in the dorsal striatum. Methamphetamine-evoked mouse locomotion behavior was recorded for 30 minutes and we selected mice with clockwise rotation behavior. To confirm the dopaminergic lesion, we conducted DAT immunostaining after all experiments ended. During the recovery time, mice were fed a high-energy diet (CMF sprout; Oriental Yeast Co., Kyoto, Japan) after surgery instead of normal chow to facilitate body weight recovery. The mice whose dopaminergic terminals were successfully lesioned were then treated with L-DOPA or saline. L-DOPA (20 mg/kg, i.p.) was administered once daily for 2 weeks in combination with benserazide hydrochloride (12 mg/kg, i.p.). All chemicals were purchased from Sigma-Aldrich (Tokyo, Japan).

### Behavioral test to evaluate LID

Open-field tests were performed to observe LID-like behaviors in the mice. The mice were placed in a 23- × 23-cm white box. A 30-minute video recording with L-DOPA treatment was conducted every day for 2 weeks. We observed the following abnormal involuntary movements: locomotive (increased locomotion with contralateral rotations), axial (contralateral dystonic posture of the neck and upper body toward the side contralateral to the lesion), and limb (jerky and fluttering movements of the limb contralateral to the side of the lesion). We counted these behaviors every 5 minutes. The mice with significant abnormal involuntary movements were used for further analyses.

### Generating TD model mice

Mice received intramuscular injections of 83 mg/kg haloperidol-decanoate (Janssen Pharmaceutical K.K., Tokyo, Japan), which releases haloperidol slowly in the hindlimb. The control mice received vehicle (sesame oil).

### Observation of VCMs by video recording

Prior to the video recording, the head frame was surgically placed on the mouse head two weeks before administration of haloperidol-decanoate. Mice were anesthetized with a mixture of ketamine and xylazine (i.p., 100 mg/kg and 10 mg/kg, respectively) and fixed in a stereotaxic apparatus. The skull surface was exposed and the periosteum and blood were removed. The head frame (12 × 19 mm, Narishige Scientific Instrument) was placed on the head and fixed with dental cement (Super-Bond C&B, Sun Medical, Shiga, Japan). The mice were allowed to recover for 1 week before starting video recording.

The mice were fixed in a stereotaxic apparatus with a holder of the head frame (MAG-1, Narishige Scientific Instrument). We zoomed in on the mouth of each mouse, and 20 minutes of mouth movements were recorded once per week for 6 weeks. After the first recording (0w, see Figure. 1C), the mice were received haloperidol-decanoate. VCMs, operationally defined as single mouth openings in the vertical plane that were not directed toward physical material, were then counted. Mice with significant VCMs were used for further analyses.

For pharmacological studies with the TD model mice, L-DOPA (20 mg/kg, i.p.) or valbenazine tosylate (0.5 and 1.5 mg/kg, p.o., Selleck Biotech, Tokyo, Japan) was administered once per day for consecutive 2 weeks. The L-DOPA administration was started two weeks after haloperidol decanoate injection and the valbenazine administration was started four weeks after the injection.

### Observation of VCMs by EMG recording

VCMs were also observed by EMG recordings. A detailed method of long-term recordings in the masseter and digastric muscles using electroencephalography EMG has been previously described ^41^. Mice were anesthetized by i.p. injection with a combination of medetomidine hydrochloride (0.75 mg/kg, Domitor®; Nippon Zenyaku Kogyo Co., Ltd., Fukushima, Japan), midazolam hydrochloride (4.0 mg/kg, Dormicum®; Sandoz K.K., Tokyo, Japan), and butorphanol tartrate (5.0 mg/kg, Vetorphale®; Meiji Seika Pharma Co., Ltd., Tokyo, Japan). A stainless-steel screw (M1-2, Unique Medical, Tokyo, Japan)—to which a urethane-coated stainless-steel wire (diameter: 0.12 mm, Unique Medical) had been attached by soldering before surgery—was implanted into the skull and used as a ground. Paired Teflon-coated stainless wires (#AS631, Cooner Wire, Chatsworth, CA, USA) were inserted into the bilateral masseter and digastric muscles. All wires were connected to a 10-pin socket used as a connector and firmly fixed to the skull with dental resin cement (56849, 3M Dental Products, St Paul, MN, USA). After surgery, atipamezole (0.75 mg/kg, Antisedan®, Nippon Zenyaku Kogyo Co., Ltd.) was administered as a medetomidine hydrochloride antagonist.

Mice were housed individually in breeding racks for at least 1 week for recovery from implantation surgery. During the subsequent 6 days, they were transported from the vivarium to the recording room, where a recording cable (TY213-042, Unique Medical) was connected and the animals were allowed to habituate to the recording conditions. *Ad libitum* access to pellets and distilled water was provided in the recording room. On the last day of habituation, a 2-hour recording session was started at 13:00 (day 0).

EMG signals were amplified (AB-611J device, NIHON KODEN Corp., Tokyo, Japan) to an optimal bandwidth (100–1000 Hz). Data obtained from EMG were digitized at 4 kHz using a PowerLab 8/35 analog-to-digital converter (PL3508, AD Instruments Inc., Dunedin, New Zealand) and stored on a personal computer with the Lab Chart 7 software package (AD Instruments). EMG activities of the masseter and digastric muscles were first rectified and then integrated for every 10-second epoch using Lab Chart 7 software.

After finishing the recording on day 0, each mouse received an intramuscular injection of haloperidol decanoate and was returned to their cage in the recording room for 6 days, with *ad libitum* access to pellets and distilled water. On days 7, 14, and 21, the mice were transferred again from their cages to the recording apparatus at 11:00; a recording session was started at 13:00 that lasted for 2 hours.

The mean integrated values for the masseter and digastric EMG activities obtained for 2 hours on day 0 were calculated for each animal; each mean value was scaled to correspond to 100%. Mean integrated values for the masseter and digastric EMG activities during the 2-hour periods on days 7, 14, and 21 in each mouse were then normalized using the values from day 0.

### Plasmid construction and AAV preparation

For the AAV vector production, we prepared the following constructs: pAAV-hSyn-ALFA-VGAT-pA, pAAV-hSyn-GAD67-1D4-pA, pAAV-hSyn-GAD65-HA-pA, and pAAV-CaMKII (0.3)-EGFP-miR30-shRNA VGAT-WPRE-pA. We synthesized ALFA (SRLEEELRRRLTE)-mouse VGAT cDNA, mouse GAD67-1D4 (TETSQVAPA) cDNA, or mouse GAD65-HA (YPYDVPDYA) cDNA, and inserted it into pAAV hSyn-pA plasmid respectively. We used previously described VGAT shRNA sequence ^42^.

AAVs were produced and their titers were measured. In brief, plasmids for the AAV vector, pHelper (Stratagene, San Diego, CA, USA) and RepCap5 (Applied Viromics, Fremont, CA, USA), were transfected into HEK293 cells (AAV293, Stratagene). After 3 days, the cells were collected and AAVs were purified twice using iodixanol. The titers for AAVs were estimated using quantitative PCR. The following AAV vectors were used in the experiments: AAV_5_-hSyn-ALFA-VGAT-pA (2×10^13^ GC/ml), AAV_5_-hSyn-GAD67-1D4-pA (2×10^13^ GC/ml), AAV_5_-hSyn-GAD65-HA-pA (2×10^13^ GC/ml), AAV_5_-CaMKII-EGFP-miR30-shRNA VGAT-WPRE-pA (3×10^13^ GC/ml), and AAV_5_-hSyn-EGFP-pA (#50465, Addgene).

### Viral vector injection

For LID and TD model mice, AAVs were injected 2 or 3 weeks before the injection of 6-OHDA or haloperidol, respectively. Mice were anesthetized with a mixture of ketamine and xylazine (i.p., 100 mg/kg and 10 mg/kg, respectively) and fixed in a stereotaxic apparatus. The skull surface was exposed and the periosteum and blood were removed. Next, 0.2 μL viral solution was injected into the right dorsal striatum (for LID) or the bilateral striatum (for TD) through a small hole (+0.6 mm anteroposterior and ±2.2 mm mediolateral from bregma and −2.2 mm dorsoventral from the brain surface according to the atlas of Paxinos and Franklin) using a glass micropipette connected to a Nanoliter 2020 injector.

### Histology

Mice were deeply anesthetized with ketamine (100 mg/kg, i.p.) and xylazine (10 mg/kg, i.p.) and perfused with a 4% paraformaldehyde phosphate buffer solution. The brains were removed from the skull and postfixed in the same fixative overnight. Subsequently, the brains were cryoprotected in 20% sucrose overnight and frozen. The frozen brains were cut on a cryostat at 25-µm thickness for ISH and mounted on silane-coated glass slides (Matsunami Glass, Osaka, Japan). For the floating sections for immunostaining, brains were cut on a cryostat at 40-µm thickness. The sections were incubated with primary antibodies overnight at room temperature. The following antibodies were used (Table S1): anti-DAT (1:1000 dilution; guinea pig polyclonal, Frontier Institute, Hokkaido, Japan), anti-VGAT (1:1000 dilution; guinea pig polyclonal, Frontier Institute); anti-PV (1:1000 dilution; goat polyclonal, Frontier Institute); anti-PLP (1:1 dilution; rat monoclonal, clone AA3 hybridoma supernatant) ^43^; anti-MAP2 (1:1000 dilution; goat polyclonal, Frontier Institute); anti-Tubb3 (1:1000 dilution; guinea pig polyclonal, Frontier Institute); anti-Bassoon (1:1000 dilution; mouse monoclonal, SAP7F407, Enzo Life Sciences, Farmingdale, NY, USA); anti-ALFA (1:500 dilution; rabbit polyclonal, NanoTag, Göttingen, Germany); anti-HA (1:500; rat monoclonal, 3F10, Roche, Basel, Switzerland); anti-rhodopsin (1:500 dilution; mouse monoclonal, 1D4, Santa Cruz Biotechnology, Dallas, TX, USA), anti-GFP (1:250 dilution, goat polyclonal; Rockland Immunochemicals, Pottstown, PA, USA), anti-NeuN (1:1000 dilution, rabbit monoclonal, EPR12763, Abcam, Cambridge, UK), anti-VGluT1 (1:1000 dilution; rabbit polyclonal, Frontier Institute), anti-VGluT2 (1:1000 dilution; rabbit polyclonal, Frontier Institute), anti-gephyrin (1:1000 dilution, mouse monoclonal, mAb7a, Synaptic Systems, Coventry, UK), anti-GLT1 (dilution; rabbit polyclonal, Frontier Institute), anti-Iba1 (dilution; rabbit polyclonal, FUJIFILM Wako, Tokyo, Japan), and anti-laminin α2 (dilution; rat monoclonal, 4H8-2; Santa Cruz Biotechnology). The sections were then treated with species-specific secondary antibodies conjugated to Alexa Fluor 488, 555, 594, or 647 for 2 hours at room temperature. Mounting medium (ProLong Glass Antifade Mountain, Thermo Fisher Scientific, Waltham, MA, USA) was applied to the samples and they were mounted with coverslips (thickness: No.1, Matsunami Glass). Macro-fluorescent images were obtained with an inverted microscope (BZ-X710; Keyence, Osaka, Japan) or a confocal microscope (LSM710; Carl Zeiss, Oberkochen, Germany). Micro-fluorescent images were obtained using SRM.

### SRM

Structured illumination microscopy (SIM)-SRM micro images were obtained using a Zeiss ELYRA 3D-SIM system equipped with an EM-CCD camera (Carl Zeiss). Before obtaining the SIM-SRM images, precise alignment for the different wavelengths of light was performed using the same mounting medium (ProLong Glass) containing 0.1% Tetraspeck (0.2-µm beads, Thermo Fisher Scientific) to correct for unavoidable laser misalignment and optical aberrations, which can lead to the alignment not coinciding at very high resolution. Next, 14–20 Z-section images were obtained at intervals of 126 nm using a 64× objective lens. The number of pattern rotations of the structured illumination was adjusted to 3 in the ELYRA system. After obtaining all images, the SIM images were reconstructed and aligned using the channel alignment data.

### Analysis of SRM images

SRM images were analyzed using ImageJ (http://rsb.info.nih.gov/ij/). Optimal brightness and grayscale pixel values were manually adjusted to provide the sharpest discrimination of the microstructure border. These adjusted images were then converted into binary images. To calculate the density of VGAT^+^, VGLuT1^+^, VGluT2^+^, Bassoon^+^, or gephyrin^+^ puncta, SRM images of each staining were randomly obtained at five locations each within the GPe and SNr. The numbers of puncta were counted and the mean number over the five locations was calculated. To calculate puncta area, 3D-SRM images of each staining were randomly obtained at three locations each within the GPe and SNr. We looked at the 3D image and used the single plane at which the area was the largest to calculate the area. We measured the area of 150 puncta from each of the three locations. The mean area of the 150 puncta was taken as the representative puncta area for each animal. To calculate the area of the PV^+^ or NeuN^+^ soma of GPe and SNr neurons, 3D-SRM images of PV or NeuN staining were randomly obtained within the GPe and SNr. We looked at the 3D image and used the single plane at which the area was the largest to calculate the area. We calculated the area of 20–30 somas per mouse and took the mean area as the representative soma area for each animal. To calculate the diameter of MAP2^+^ dendrites of GPe and SNr neurons, 3D-SRM images of MAP2 staining were randomly obtained within the GPe and SNr. The diameters of 40–50 MAP2^+^ dendrites per mouse were measured and the mean diameter was taken as the representative dendrite diameter for each animal. Methods used to measure the axon diameter, myelin thickness, and g-ratio (the ratio of the inner axonal diameter to the total outer diameter) have been previously described ^44, 45^. Axon diameter was defined as the minor axis of an ellipse-approximated axon. The median axon diameter over 50 myelinated axons was considered the representative axon diameter value in each animal. The g-ratio was calculated using the equation 0.5AD / (0.5AD + MT), where AD is the axon diameter and MT is myelin thickness. The mean myelin thickness and g-ratio over 50 myelinated axons were considered the representative myelin thickness and g-ratio, respectively, for each animal. To calculate the percentage area of GLT1^+^ astrocytes and Iba1^+^ microglia, confocal images of GLT1 or Iba1 staining were randomly obtained at five locations each within the GPe and SNr. The percentage area was calculated using the equation: grail area / image area × 100, where grail area is the GLT1^+^ or Iba1^+^ area and image area is the area of the ROI. The average area of five locations was taken as the representative percentage area of astrocytes or microglia for each animal.

### ISH

The detailed protocol has been described previously ^44^. The 25-µm sections were treated with 40 μg/mL proteinase K (Roche) for 30 minutes before being washed with phosphate-buffered saline (PBS) for 5 minutes and postfixed with 4% paraformaldehyde in PBS for 15 minutes to inactivate the proteinase K. After a 5-minute wash with PBS, the sections were acetylated with 0.25% acetic anhydride. Prehybridization was conducted for 2 hours at 60°C in prehybridization buffer containing 50% formamide (Wako, Tokyo, Japan), 50× Denhardt’s solution (Nacalai Tesque, Kyoto, Japan), and 10 mg/mL salmon sperm DNA (Invitrogen, Tokyo, Japan). After removing the prehybridization buffer, sections were hybridized overnight at 60°C in hybridization buffer containing the following digoxigenin-labeled cRNA probes (Table S1): *Slc32a1* (VGAT, nt_241-2805, NM_009508), *Gad1* (GAD67) ^46^, *Gad2* (GAD65) ^47^, colony-stimulating factor 1 receptor (*Csf1r*, a marker of microglia) ^48^, *Plp1* (an oligodendrocyte marker) ^49^, *Gja1* (connexin 43, an astrocyte marker, nt_33-3097, NM_010288), and *Pdgfra* (an OPC marker) ^48^. After the sections were washed in buffers with serial differences in stringency, they were incubated with an alkaline phosphatase-conjugated anti-digoxigenin antibody (1:5000; Roche) for 90 minutes at room temperature. Unbound antibody was removed by four 10-minute washes. The cRNA probes were visualized by being incubated with a freshly prepared colorimetric substrate (NBT/BCIP; Roche) overnight at room temperature. Following ISH staining, the sections were counterstained with nuclear fast red (Sigma-Aldrich) and images were captured using an inverted light microscope (BZ-X710, Keyence).

### Cell counts of ISH images

To evaluate the cell density of astrocytes, microglia, OPCs, and oligodendrocytes, cell counts were performed from the ISH images using ImageJ. Cell density was calculated by dividing the number of cells expressing the marker RNA in an ROI by the area of the ROI. The ROI was defined as a region that included the GPe and SNr.

### qRT-PCR

Mice were sacrificed by cervical dislocation and CPu, GPe, and SNr tissue was collected. Total RNA was isolated from these regions using TRIzol (Thermo Fisher Scientific) and reverse transcribed into cDNA using ReverTra Ace qPCR RT Master Mix (Toyobo, Osaka, Japan). The qRT-PCR was performed using TaqMan probes (Thermo Fisher Scientific) on the Step One Plus real-time PCR monitoring system (Thermo Fisher Scientific). The primers and TaqMan probe sequences of *Slc32a1* (VGAT), *Gad2* (GAD65), and *Gad1* (GAD67) were Mm00494138_m1, Mm00484623_m1, and Mm00725661_s1, respectively. The expression of these mRNA transcripts was measured; mRNA levels were normalized to those of glyceraldehyde 3-phosphate dehydrogenase (*Gapdh*, Mm99999915_g1).

### MRI

An *ex vivo* MRI study of the LID model mice was performed using a 9.4 T BioSpec 94/30 (Biospin GmbH, Ettlingen, Germany) unit and a solenoid-type coil with 28-mm inner diameter for transmitting and receiving. After the final L-DOPA administration, mice were deeply anesthetized with ketamine (100 mg/kg) and xylazine (10 mg/kg) and perfused with a 4% paraformaldehyde phosphate buffer solution. The brains were removed with the skull and postfixed in the same fixative for 24 hours. The fixed brains were then stored in PBS for 1 week. The duration of paraformaldehyde and PBS immersion was the same for all samples to avoid differences in brain volume caused by postperfusion immersion fixation and storage (Guzman et al., 2016). Four brains (with their skulls) were firmly fixed using fitted sponges into an acrylic tube (30-mm diameter) filled with Fluorinert (Sumitomo 3M Limited, Tokyo, Japan) to minimize the signal intensity attributed to the embedding medium. Additionally, vacuum degassing was performed to reduce air bubble-derived artifacts of structural images. For volume analysis, structural images were acquired using T2-weighted multi-slice rapid acquisition with relaxation enhancement (RARE) with the following parameters: repetition time = 20,000 ms, echo time = 15 ms, spatial resolution = 100 × 100 × 100 µm, RARE factor = 4, and 24 averages.

Brain volumes of LID model mice were compared between the ipsilateral and contralateral sides using ROI-based analysis. For each hemisphere, ROIs of 64 brain loci (128 loci in total) were defined using the Allen Brain Atlas-based flexible annotation atlas of the mouse brain (Table S1) ^50^. When we compared the voxel size of each ROI, there were small differences in voxel size between hemispheres, indicating a risk of false positives for brain volume changes. We therefore chose ROI-based analysis for the brain volume analysis (instead of voxel-based analysis). We made tissue probability maps (TPMs) of gray matter, white matter, and cerebrospinal fluid from the flexible annotation atlas for the preprocessing of tissue segmentation. The GPe and GPi of the TPMs that were commonly used were classified as white matter. We thus made the TPMs where GPe and GPi were classified as gray matter.

Preprocessing and statistics for ROI-based brain volume analysis were performed using SPM12 (Wellcome Trust Centre for Neuroimaging, London, UK) and in-house software written in MATLAB (MathWorks, Natick, MA, USA). First, each T2-weighted image was resized by a factor of 10 to account for the whole-brain volume difference between humans and rodents. It was then re-sampled into 1-mm isotropic voxels and aligned to the same space by registering each image to the TPM. Next, each image was segmented into TPMs of gray matter, white matter, and cerebrospinal fluid using the unified segmentation approach, which enables image registration, tissue classification, and bias correction. The segmented images were spatially normalized into the population template, which was created by diffeomorphic anatomical registration through the Diffeomorphic Anatomical Registration Through Exponentiated Lie Algebra (DARTEL) algorithm. Modulated gray matter images were then obtained for each animal by the determinant of the Jacobian of the transformation to account for the expansion and/or contraction of brain regions. These images were smoothed with a 3-mm full width at half maximum Gaussian kernel. The modulated values referring to brain volumes were extracted from each ROI and averaged within each ROI. Two-tailed two-sample t-tests were performed to compare the averaged values at each ROI between the ipsilateral and contralateral sides. The p-values were corrected using false discovery rate correction.

### EM

Mice were deeply anesthetized with ketamine (100 mg/kg, i.p.) and xylazine (10 mg/kg, i.p.) and perfused with 4% paraformaldehyde and 2.5% glutaraldehyde in 0.1 M phosphate buffer (pH 7.4). The brains were removed and postfixed in the same fixative overnight before being cut at 1-mm thickness on a vibratome. Slices were then treated with 2% OsO_4_ (Nisshin EM, Tokyo, Japan) in 0.1 M cacodylate buffer containing 0.15% K_4_[Fe(CN)_6_] (Nacalai Tesque), washed four times with cacodylate buffer, and incubated with 0.1% thiocarbohydrazide (Sigma-Aldrich) for 20Lminutes and 2% OsO_4_ for 30Lminutes at room temperature. The slices were then treated with 2% uranyl acetate at 4°C overnight and stained with Walton’s lead aspartate at 50°C for 2Lhours. The slices were dehydrated through a graded ethanol series (60%, 80%, 90%, 95%, and 100%) at 4°C; infiltrated sequentially with acetone dehydrated with a molecular sieve, a 1:1 mixture of resin and acetone, and 100% resin; and embedded with Aclar film (Nisshin EM) in Durcupan resin with carbon (Ketjen black) (Sigma-Aldrich). The specimen-embedded resin was polymerized at 40°C for 6 hours, 50°C for 12 hours, 60°C for 24 hours, and 70°C for 2 days. After trimming the region containing the GPe from the brain, the samples were serially imaged with a Merlin (Carl Zeiss) electron microscope equipped with the 3View system and an OnPoint backscattered electron detector (Gatan, Pleasanton, CA, USA). The Merlin is a field emission-type scanning electron microscope with a single electron beam, which was set to 1.2–1.5 kV acceleration voltage and 130 pA beam current.

For EM image analysis, serial images of the serial block face scanning EM were handled with Fiji/ImageJ and segmented using Microscopy Image Browser (http://mib.helsinki.fi/). The EM images were acquired in the ipsilateral GPe of the LID model mouse and its control mouse. The area of terminals making inhibitory synaptic contacts with dendrites or somas, the diameter of dendrites surrounded by the terminal, and the diameter of unmyelinated axons were measured. Inhibitory terminals were defined as those with vesicles present and symmetrical postsynaptic density.

### Mass spectrometry imaging of GABA and dopamine

Mice were deeply anesthetized with ketamine (100 mg/kg, i.p.) and xylazine (10 mg/kg, i.p.), and the brains were rapidly removed from the skull and frozen in liquid N_2_. Next, 10-μm-thick sections of fresh-frozen brains were prepared with a cryostat and thaw-mounted on conductive indium-tin-oxide-coated glass slides (Matsunami Glass).

A pyrylium-based derivatization method was applied for the tissue localization imaging of neurotransmitters ^51, 52^. TMPy solution (4.8 mg/200 μL; Taiyo Nippon Sanso Co., Tokyo, Japan) was applied to brain sections using an airbrush (Procon Boy FWA Platinum 0.2-mm caliber airbrush, Mr. Hobby, Tokyo, Japan). To enhance the reaction efficiency of TMPy, TMPy-sprayed sections were placed into a dedicated container and allowed to react at 60°C for 10 minutes. The container contained two channels in the central partition, to wick moisture from the wet filter paper region to the sample section region. The filter paper was soaked with 1 mL methanol/water (70/30 volume/volume) and placed next to the section inside the container, which was then completely sealed to maintain humidity levels. The TMPy-labeled brain sections were sprayed with matrix (CHCA-methanol/water/TFA = 70/29.9/0.1 volume/volume) using an automated pneumatic sprayer (TM-Sprayer, HTX Tech., Chapel Hill, NC, USA). Ten passes were sprayed according to the following conditions: flow rate, 120 µL/min; airflow, 10 psi; nozzle speed, 1100 mm/minute.

To detect the laser spot area, the sections were scanned and laser spot areas (200 shots) were detected with a spot-to-spot center distance of 80 µm. Signals between *m*/*z* 100–650 were corrected. The section surface was irradiated with yttrium aluminum garnet laser shots in the positive ion detection mode using matrix-assisted laser desorption/ionization time-of-flight mass spectrometry (MALDI-TOF MS; timsTOF fleX, Bruker Daltonics, Bremen, Germany). The laser power was optimized to minimize the in-source decay of targets. Obtained mass spectrometry spectra were reconstructed to produce mass spectrometry images using Scils Lab software (Bruker Daltonics). Optical images of brain sections were obtained using a scanner (GT-X830, Epson, Tokyo, Japan) followed by MALDI-TOF MS of the sections. The detected masses of TMPy-labeled standard GABA (*m*/z 208.163) increased by 105.0 Da compared with the original mass (molecular weight 103.0 Da). Tandem mass spectrometry confirmed the fragmentation ions of TMPy from the standard sample. A fragmented ion of the pyridine ring moiety (*m/z* 122.1) was regularly cleaved and observed for all TMPy-modified target molecules.

### Neuronal activity recording

Single-unit recording was performed before and after the administration of haloperidol-decanoate for TD or AAV for striatal VGAT overexpression, respectively. The surgical operation to mount a head holder onto the head of each mouse was performed as described previously (Chiken et al., 2015; Sano et al., 2013; Wahyu et al., 2021). Each mouse was anesthetized with ketamine (100 mg/kg, i.p.) and xylazine (5 mg/kg, i.p.) and held in a stereotaxic apparatus (SR-6M, Narishige Scientific Instrument). The skull was widely exposed and covered with bone adhesive resin (ESTECEM II, Tokuyama Dental, Tokyo, Japan). A small U-shaped head holder made of acetal resin was attached to the skull with acrylic resin (Unifast II, GC, Tokyo, Japan). The mouse was thus held in the stereotaxic apparatus with its head restrained using the U-shaped head holder. For the TD model mice, part of the skull over the right hemisphere was removed to access the motor Cx, CPu, GPe, and SNr. Somatotopy of the motor Cx was confirmed by intracortical microstimulation (a train of 10 pulses at 333 Hz, 200-µs duration, <20 µA). Two pairs of bipolar stimulating electrodes (50-µm diameter Teflon-coated tungsten wires, tip distance 300–400 µm) were chronically implanted into the orofacial and forelimb regions of the primary motor Cx and fixed using acrylic resin ^23, 53, 54^. For mice who received AAV injections, part of the skull over the right hemisphere (ipsilateral to the AAV injection) was removed to access the motor Cx, GPe, and SNr. Somatotopy of these regions was confirmed by intracortical microstimulation (a train of 10 pulses at 333 Hz, 200-µs duration, <20 µA).

After recovering from the surgery, each awake mouse was positioned painlessly in a stereotaxic apparatus (SR-6M) using the U-shaped head holder ^23, 53, 54^. For single-unit recording, a glass-coated tungsten microelectrode (0.5 or 1.0 MΩ at 1 kHz; Alpha Omega, Nazareth, Israel) was inserted vertically into the right GPe (target area: posterior 0.3–0.6 mm and lateral 2.2–2.6 mm from bregma) or SNr (posterior 2.6–3.0 mm and lateral 1.6–2.2 mm from bregma) through the dura mater using a hydraulic microdrive. Signals from the microelectrode were amplified and filtered (0.3–5.0 kHz). Unit activity was isolated, converted to digital data with a homemade time-amplitude window discriminator, and sampled at 2.0 kHz using a computer with LabVIEW 2013 software (National Instruments, Austin, TX, USA) for offline data analysis. The responses to Cx electrical stimulation (200-µs duration, single monophasic pulse at 0.7 Hz, 50-µA strength) through the stimulating electrodes implanted in the motor Cx were examined by constructing PSTHs (bin width, 1 ms; prestimulus, 100 ms; poststimulus, 800 ms) for 100 stimulation trials. These Cx-evoked responses were recorded in the GPe and SNr and were considered control data.

After refining the recording in control conditions, haloperidol-decanoate or AAV vector was injected. For the TD experiments, haloperidol-decanoate was injected in the hindlimb (83 mg/kg, intramuscular) as described in the *Generating TD model mice* section. Recordings from the GPe and SNr in TD conditions were started 3 weeks after the injection. For VGAT overexpression, AAV-hSyn-ALFA-VGAT was injected into the right striatum, ipsilateral to the recording side. Neuronal responses to Cx stimulation were examined. The striatal region with motor cortical inputs from the orofacial and/or forelimb regions was identified and AAV vector was injected (0.3 µL/site, 1 site). Approximately 3–4 weeks after the injection, when VGAT was overexpressed, recording from the GPe and SNr in VGAT overexpression conditions was started.

Responses to Cx stimulation were analyzed using PSTHs. Cx stimulation typically induced a triphasic response—composed of early excitation, inhibition, and late excitation—in GPe and SNr neurons. The mean value (μ_baseline_) and standard deviation (SD_baseline_) of the discharge rate during 100 ms preceding the onset of stimulation were considered the baseline discharge rate; the significance level was set as μ_baseline_ ± 1.65 SD_baseline_ (corresponding to p=0.1, two-tailed t-test). If at least two consecutive bins (2 ms) exceeded the significance level, the response was judged significant as described previously ^53, 54^. The initial point was determined as the time of the first bin exceeding the significance level. The responses were judged to end when two consecutive bins fell below the significance level. The endpoint was determined as the time of the last bin exceeding this level. The duration (from the initial point to the endpoint) and the amplitude (the area of response; the number of spikes during the significant changes minus the number of spikes during baseline) of each response were calculated and compared ^53, 54^. If there was no significant early excitation, inhibition, or late excitation, the duration and amplitude were set to zero. For averaged PSTHs, the PSTH of each neuron with a significant response to cortical stimulation was averaged within the same conditions and smoothed using a binomial filter (σ = 2.0 ms).

### Forelimb reaching task

Detailed methods are provided in a previous study ^28^. Before 6-OHDA injections, mice were food-restricted and trained. Their body weights were maintained at 85% of their initial body weight. The training chamber was constructed as a clear Plexiglas box (20 cm tall, 15 cm deep, and 8.5 cm wide) into which each mouse was placed. There was one vertical slit (0.5 cm wide and 13 cm high) in the center of the front wall of the box. A 1.25-cm-tall exterior shelf was affixed to the wall in front of the slits to hold food pellets (10 mg each: Dustless Precision Pellets, Bio-Serv, Prospect, CT, USA) for a food reward. The training period (1–5 days in duration) was used to familiarize mice with the training chamber and task requirements and to determine their preferred limbs. Food pellets were placed in front of the center slit, and mice used both paws to reach for them. The training was finished when 50 reach attempts were achieved within 30 minutes and the mouse showed 70% limb preference. After training, the pre-period reaching task was conducted for 7 days; each day consisted of one session of 100 trials with the preferred limb or 30 minutes. Food pellets were presented individually in front of the slit. Next, 6-OHDA was injected into the medial forebrain bundle contralateral to the preferred limb. Two days after the injection, the 6-OHDA-period reaching task (after injection of 6-OHDA) was conducted for 6 days. Mice then underwent surgery to implant one L-DOPA pellet (hormone and drug pellets of the matrix-driven delivery system; 15 mg/pellet, Innovative Research of America, Sarasota, FL, USA) to continuously administer L-DOPA, and the L-DOPA-period reaching task (after the implantation of the L-DOPA pellet) was conducted for 14 days.

To evaluate the task, mice displayed three reach attempt types: fail, drop, or success. A fail was scored as a reach in which the mouse failed to touch the food pellet or knocked it away. A drop was a reach in which the mouse retrieved the food pellet but dropped it before putting it into its mouth. A success was a reach in which the mouse retrieved the pellet and put it into its mouth. Occasionally a mouse used the non-preferred limb; this was categorized as a fail. Success rates were calculated as the percentage of successful reaches relative to the total reach attempts.

### Statistical analysis

Statistical processing was performed using MATLAB and Excel (Microsoft, Redmond, WA, USA) software. Paired t-tests were performed to compare the contralateral and ipsilateral sides. Two-tailed Student’s t-tests were performed to compare TD and its control mice. Two-way repeated analysis of variance (ANOVA) was performed to compare the dyskinesia time course of LID mice. Bonferroni corrections were applied to correct p-values for multiple comparisons. Values are shown as the mean and standard error of the mean (SEM), and are plotted as scatter diagrams.

