## Supplemental information for "Shared GABA transmission pathology in dopamine agonist- and antagonist-induced dyskinesia"

**Supplementary text (related to Figure 1)**

**Anatomical signature of MSN-terminating nuclei**

Striatopallidal and striatonigral MSNs terminate in the GPe and SNr, respectively. The axons of both MSN populations are unmyelinated ^52^ and have VGAT^+^ presynaptic terminals. Regardless of MSN type, MSN target nuclei share a similar structure; they consist of gray matter with fragmented white matter bundles composed of the myelinated axons of cortical pyramidal neurons (Figure S1G). SRM of GPe gray matter demonstrated that VGAT^+^ puncta surrounded PV^+^ dendrites and soma of GPe principal cells (Figure 1F). EM demonstrated vesicle-rich presynaptic terminals surrounding dendrites and soma, and thin, unmyelinated axons occupying the neuropil of GPe gray matter (Figure 1F).

**Increased gray and white matter volumes in** the **GPe/SNr of LID mice**

To address whether gray or white matter volume increases contributed to the GPe/SNr volume increase in the LID model, we conducted PLP immunohistochemistry and quantified the areas of gray (PLP^−^, neuropil area) and white (PLP^+^, myelinated axon area) matter in ipsilateral and contralateral nuclei (Figure S1I). In the ipsilateral GPe and SNr, both gray and white matter areas were larger than those in the contralateral hemisphere (Figure S1J). The ipsilateral proportion of gray matter was larger than that of the contralateral hemisphere, indicating that increased gray matter volume contributes more to the total nucleus size (Figure S1K).

**Increased sizes of presynaptic and postsynaptic structures in the GPe/SNr of LID mice**

To examine the neuronal elements that contributed to increased gray matter volume in the GPe and SNr, we labeled presynaptic MSN terminals, unmyelinated MSN axons, and the soma and dendrites of principal neurons using immunohistochemistry. Signals were detected using SRM. The size and density of VGAT^+^ puncta were significantly increased in the ipsilateral GPe and SNr compared with those in the contralateral hemisphere (Figure 1G), indicating that both the volume and number of MSN presynaptic terminals is increased in LID. In contrast, these features were comparable in control mice (Figure S2A). The EM analyses strengthened the SRM findings; presynaptic terminals that were associated with a dendrite increased in size in LID but not control mice (Figure S2B, C). Consistent with the increased density of MSN terminals, the percentage area of unmyelinated axons (βIII tubulin [Tubb3]^+^, microtubule-associated protein 2 [MAP2]^−^, and PLP^−^) was increased in the ipsilateral GPe and SNr (Figure S2E, F).

Principal neurons in the GPe and SNr are divided into PV^+^ and PV^−^ populations ^53^. We conducted NeuN and MAP2 immunohistochemistry to identify soma and dendrites, respectively, and evaluated their sizes in each population. The soma areas of PV^+^ neurons were significantly increased in the ipsilateral GPe and SNr, whereas those of PV^−^ neurons were significantly increased in the GPe but not the SNr (Figure 1G and Figure S2H). The dendrite diameters of PV^+^ and PV^−^ neurons were significantly increased in the ipsilateral GPe (Figure S2I); this finding was confirmed by MAP2 staining (Figure S2G) and EM analyses (Figure S2D). In addition, we conducted VGAT and gephyrin (a postsynaptic maker of inhibitory synapses) immunohistochemistry to examine the sizes of VGAT^+^ and gephyrin^+^ puncta. Their sizes were significantly increased in the ipsilateral GPe and SNr and were positively correlated (Figure S2J, K). These results indicate that enlargement of the soma and dendrites of principal neurons contributes to the increased GPe/SNr volume.

Principal neurons in the GPe/SNr receive glutamatergic inputs from the cortex and STN; presynaptic terminals have vesicular glutamate transporter (VGluT) 1 and 2, respectively ^54,55^. In LID model mice, the density and area of VGluT1^+^ puncta were comparable between the GPe and SNr (Figure S3A). In contrast, the density of VGluT2^+^ puncta was significantly decreased and the area was increased in the ipsilateral GPe and SNr (Figure S3B). Nonetheless, the density of VGluT^+^ puncta was less than one-tenth that of VGAT^+^ puncta; thus, volume changes in glutamatergic terminals may have a negligible impact on GPe/SNr volume.

Regarding the observed white matter increase in the ipsilateral GPe and SNr, we evaluated myelinated axon diameter and myelin thickness using PLP staining and SRM ^56,57^. Both indices were significantly increased in the ipsilateral GPe and SNr (Figure S3C) and likely account for the increased white matter volume.

**Comprehensive histological analyses emphasize MSN terminal changes in LID mice**

To fully address factors that might explain GPe/SNr volume increases, we quantified the numbers (density) and volumes (percentage area) of cells including neurons, astrocytes, oligodendrocytes, oligodendrocyte precursor cells (OPCs), microglia, and vascular cells. The number of principal neurons (NeuN^+^) per nucleus was unchanged; however, the density was decreased in the ipsilateral GPe and SNr (Figure S3D), indicating an increased-volume-associated reduction in cell density and no neuronal proliferation. The number of glial cells was determined after *in situ* hybridization (ISH) with *Gja1* (astrocyte marker), *Plp1* (oligodendrocyte marker), *Pdgfra* (OPC marker), or *Csf1r* (microglial marker). GPe astrocytes (Figure S3E) and GPe/SNr oligodendrocytes (Figure S3F) showed a neuron-like pattern of cell number change, with an increased-volume-associated reduction in cell density. SNr astrocytes (Figure S3E) and GPe OPCs (Figure S3G) exhibited increased cell number with sustained cell density, indicating an adaptive response to GPe/SNr volume increase. SNr OPCs (Figure S3G) and GPe/SNr microglia (Figure S3H) showed increases in both number and density, which is likely relevant to the LID volume increase. Immunohistochemistry for glutamate transporter 1 (GLT1; astrocyte marker, labels membranes) and ionized calcium binding adapter molecule 1 (Iba1; microglia marker, labels cytoplasm) demonstrated that the percentage area of astrocytic processes was comparable between contralateral and ipsilateral GPe and SNr, whereas that of microglia was increased in the ipsilateral GPe or SNr, consistent with the cell density data (Figure S3J, K). Immunohistochemistry of laminin α2 (vasculature marker) showed an increased vasculature area in the ipsilateral GPe or SNr; however, the vasculature areas normalized by VGAT areas were comparable between contralateral and ipsilateral GPe and SNr (Figure S3I).

These results are summarized in Table S3 and indicate that there are significant increases in SNr OPCs and GPe/SNr microglia in LID. However, the populations of these cells are much smaller than those of neurons and astrocytes within the GPe and SNr ^53^. We therefore suppose that OPC-/microglia-mediated cell volume changes contributed relatively little to regional volume increases. Increases in total cell number without increased cell densities (e.g., in astrocytes and vasculature) may adaptively support an LID-induced nuclear volume increase. Together, these findings suggest that inhibitory presynaptic structures (MSN terminals) and postsynaptic structures (dendrites and soma of GPe/SNr principal neurons) as well as cortical myelinated axons are the main contributing factors to increased GPe/SNr volumes in LID.

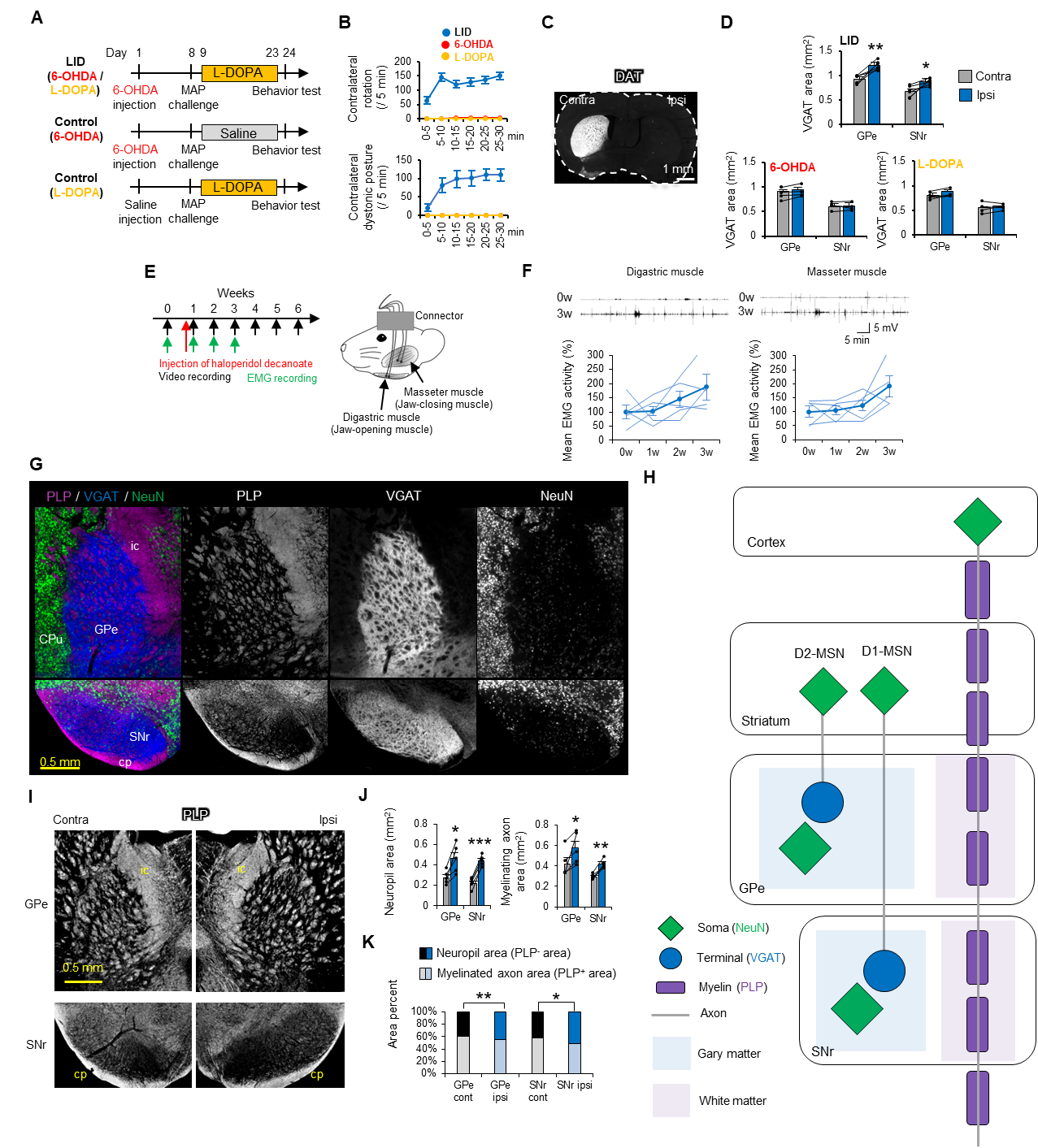

**Figure S1. Validation of LID and TD model mice and typical structure of the GPe and SNr, related Figure 1.**

(A) Time course of the generation of LID model and control mice.

(B) Number of contralateral rotations and contralateral dystonic postures were counted every 5 minutes in LID model (n=6) and control (n=6) mice.

(C) Dopamine transporter (DAT) staining confirmed dopamine depletion in the Ipsi hemisphere (relative to 6-OHDA injection) compared with the findings in the Contra hemisphere.

(D) VGAT^+^ area was compared between the Contra and Ipsi hemispheres of LID (n=4) and control (n=4) mice. (E) Time course of TD model mouse generation. VCMs were recorded by video or EMG every week. EMG was recorded from digastric (jaw-opening) and masseter (jaw-closing) muscles for 3 weeks.

(F) Representative EMG responses from the two muscles at 0 and 3 weeks. Mean EMG activity was plotted every week (n=5).

(G) Representative low-magnification images of PLP, VGAT, and NeuN in the GPe and SNr of control mice. ic; internal capsule, cp; cerebral peduncle. GPe and SNr share a similar structure. They consist of gray matter (VGAT and NeuN) with fragmented white matter bundles (PLP) composed of the myelinated axons of cortical pyramidal neurons.

(H) Schematic diagram of the anatomy of striatopallidal and striatonigral MSN-terminating nuclei.

(I) Representative images of PLP immunohistochemistry in the Contra and Ipsi GPe and SNr of LID model mice.

(J) The neuropil area (PLP^−^ area) and myelinated axon area (PLP^+^ area) were plotted for the GPe and SNr of LID model mice (n=5).

(K) Neuropil and myelinated axon areas were compared between the Contra and Ipsi GPe and SNr in LID model mice (n=5).

*p<0.05. **p<0.01, ***p<0.001 (paired t-test, p-values corrected by Bonferroni correction). Values are plotted as the mean ± SEM.

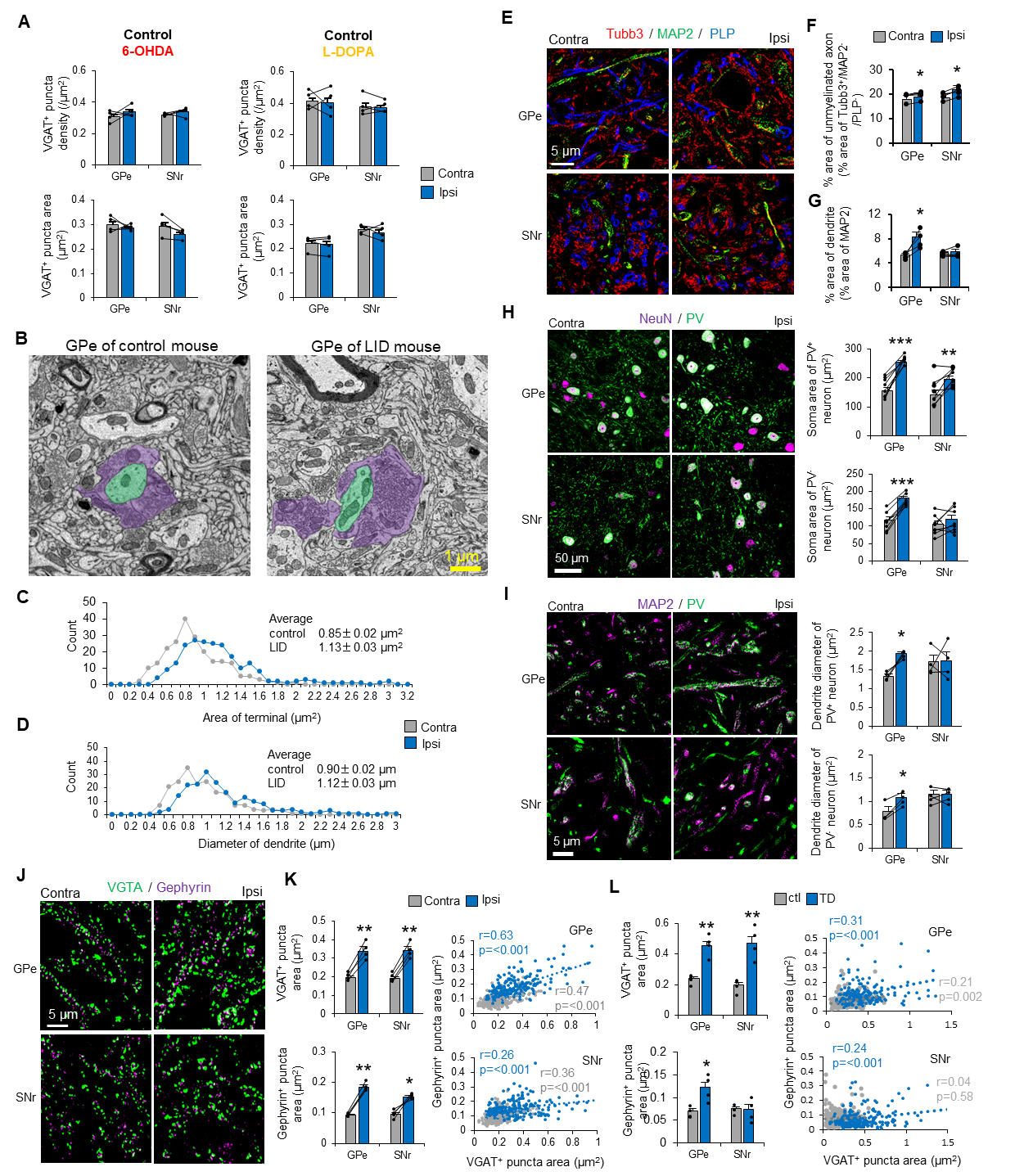

**Figure S2. The enlargement of inhibitory presynaptic terminal of the MSNs and postsynaptic soma/dendrite of GPe/SNr neurons, related to Figure 1.**

(A) The density and area of VGAT^+^ puncta of MSN terminals were compared between the Contra and Ipsi GPe and SNr in control mice for 6-OHDA (n=4) and L-DOPA (n=4) injections.

(B) Representative EM images of the GPe in control (left panel) and LID (right panel) mice. Green indicates dendrites of GPe neurons and purple indicates MSN terminals.

(C, D) Histograms of the terminal area of MSNs (C) and dendrite diameters (D) of GPe neurons compared between control (n=1) and LID (n=1) mice. For each group, 250 terminals and 200 dendrites were counted from one animal.

(E) Representative SRM images of Tubb3 (myelinated/unmyelinated axons and dendrites), MAP2 (dendrites), and PLP (myelinated axons) immunohistochemistry in LID mice.

(F) Percentage areas of unmyelinated axons (area of Tubb3/area of MAP2 and PLP) in the GPe and SNr were compared between Contra and Ipsi in LID mice (n=4).

(G) The MAP2 area was defined as the area of dendrites of GPe or SNr neurons; percentage areas were compared between Contra and Ipsi hemispheres in LID mice (n=4).

(H) Representative confocal microscopy images of NeuN and PV immunohistochemistry in LID mice. NeuN^+^ soma areas of PV^+^ and PV^−^ neurons were compared between the Contra and ipsi GPe and SNr in LID mice (n=8).

(I) Representative SRM images of MAP2 and PV immunohistochemistry in LID mice. MAP^+^ dendrite diameters of PV^+^ and PV^−^ neurons were compared between the Contra and ipsi GPe and SNr in LID mice (n=4).

(J) Representative SRM images of VGAT and gephyrin immunohistochemistry in LID mice.

(K) Areas of VGAT^+^ and gephyrin^+^ puncta of MSN terminals were compared between the Contra and ipsi GPe and SNr in LID mice (n=4). Areas of VGAT^+^/gephyrin^+^ puncta in the GPe and SNr in LID mice were plotted. More than 200 puncta were counted from four mice.

(L) Areas of VGAT^+^ and gephyrin^+^ puncta in the GPe and SNr were compared between control (n=4) and TD (n=4) mice. Areas of VGAT^+^/gephyrin^+^ puncta in the GPe and SNr were plotted for control and TD mice. More than 200 puncta were counted from four mice in each group.

*p<0.05, **p<0.01. (Student’s or paired t-test, p-values corrected by Bonferroni correction). Values are plotted as the mean ± SEM.

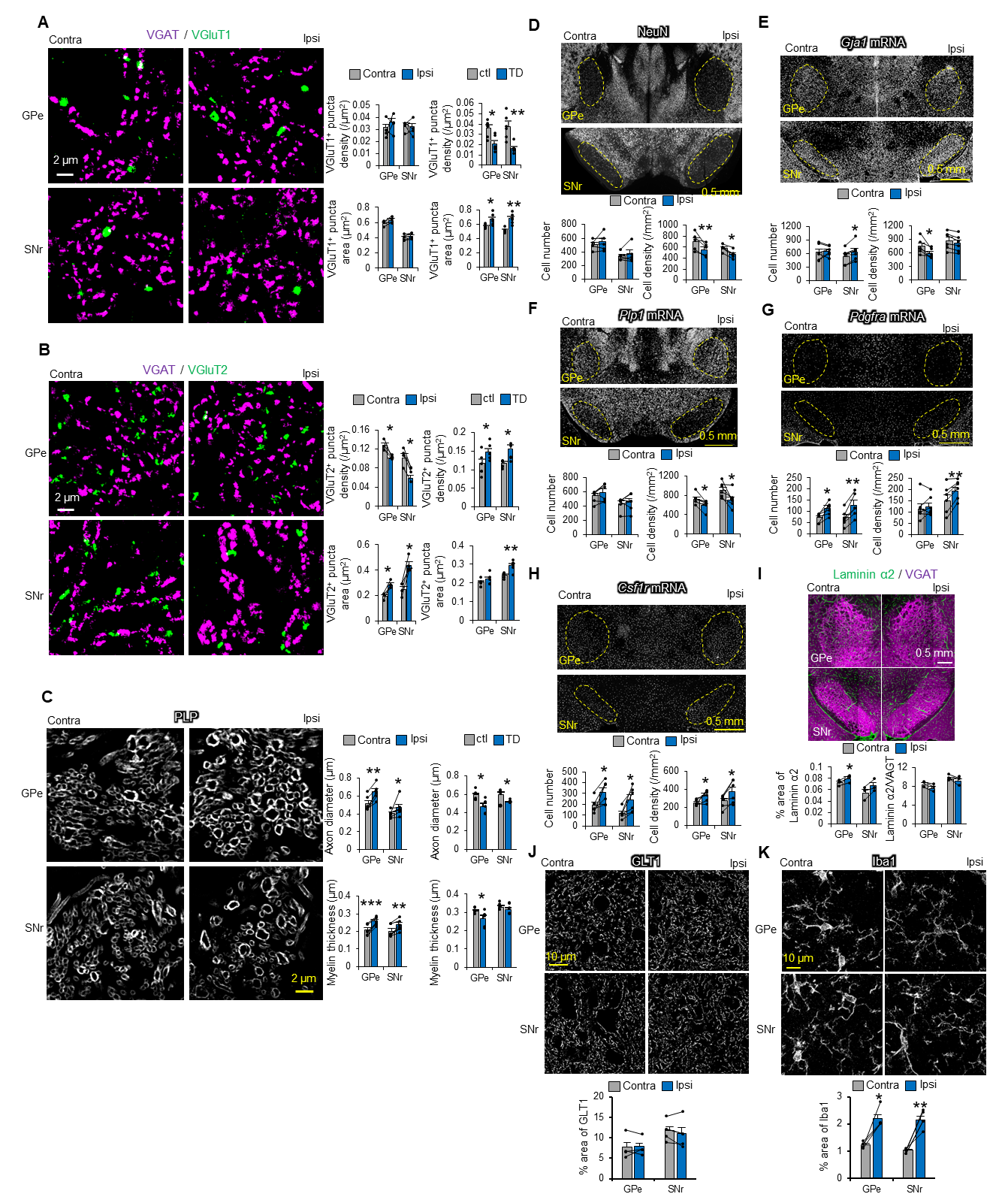

**Figure S3. Contribution of another neuronal compartments and glial cells, related to Figure 1.**

(A) Representative SRM images of VGAT and VGluT1 immunohistochemistry in LID mice. The density and area of VGluT1^+^ puncta originating from the cortex were compared between the Contra and Ipsi GPe and SNr in LID mice (n=4). They were compared between TD (n=5) and control (n=5) mice.

(B) Representative SRM images of VGAT and VGluT2 immunohistochemistry in LID mice. The density and area of VGluT2^+^ puncta originating from the STN were compared between the Contra and Ipsi GPe and SNr in LID mice (n=4). They were compared between TD (n=5) and control (n=5) mice.

(C) Representative SRM images of PLP immunohistochemistry in LID mice. Axon diameter and myelin thickness were compared between the Contra and Ipsi GPe and SNr in LID mice (n=5). They were compared between TD (n=5) and control (n=5) mice.

(D) Representative images of NeuN immunohistochemistry in LID mice. The number and density of NeuN^+^ neurons were compared between the Contra and Ipsi GPe and SNr in LID mice (n=5).

(E) Representative ISH images of *Gja1* mRNA in LID mice. The number and density of *Gja1*^+^ astrocytes were compared between the Contra and Ipsi GPe and SNr in LID mice (n=5).

(F) Representative ISH images of *Plp1* mRNA in LID mice. The number and density of *Plp1*^+^ oligodendrocytes were compared between the Contra and Ipsi GPe and SNr in LID mice (n=5).

(G) Representative ISH images of *Pdgfra* mRNA in LID mice. The number and density of *Pdgfra*^+^ OPC were compared between the Contra and Ipsi GPe and SNr in LID mice (n=5).

(H) Representative ISH images of *Csfr1r* mRNA in LID mice. The number and density of *Csfr1r* ^+^ microglia were compared between the Contra and Ipsi GPe and SNr in LID mice (n=5).

(I) Representative images of laminin α2 and VGAT immunohistochemistry in LID mice. The percentage area of laminin α2^+^ blood vessels and normalized laminin α2 area by VGAT area were compared between the Contra and Ipsi GPe and SNr in LID mice (n=3).

(J) Representative SRM images of GLT1 immunohistochemistry in LID mice. The percentage area of GLT1^+^ astrocytes was compared between the Contra and Ipsi GPe and SNr in LID mice (n=4).

(K) Representative SRM images of Iba1 immunohistochemistry in LID mice. The percentage area of Iba1^+^ microglia was compared between the Contra and Ipsi GPe and SNr in LID mice (n=4).

*p<0.05. **p<0.01, ***p<0.001 (Student’s or paired t-test, p-values corrected by Bonferroni correction). Values are plotted as the mean ± SEM.

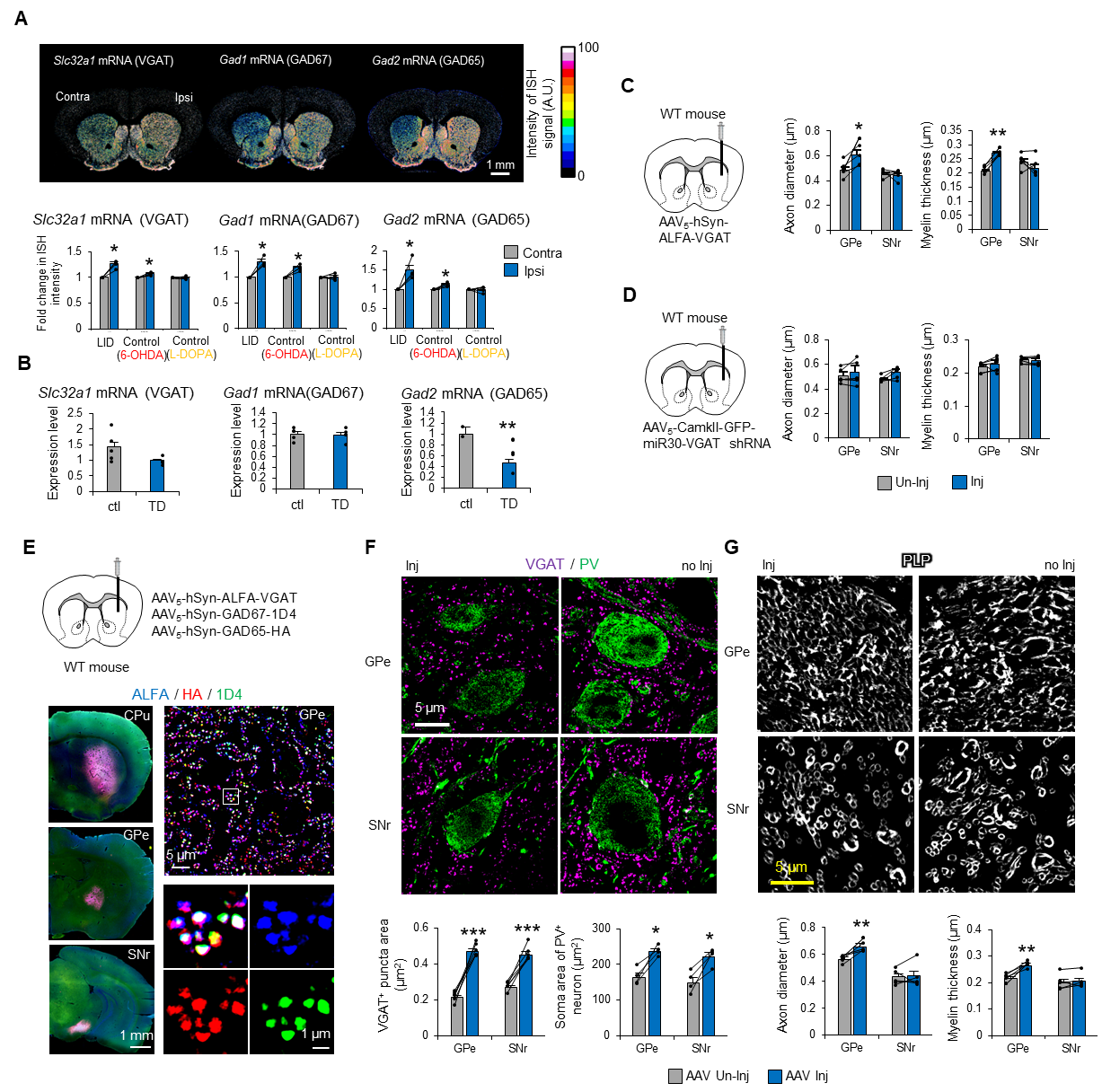

**Figure S4. Striatal overexpression of GABA-related genes was associated with dyskinesia-related pathology, related to Figure 2.**

(A) ISH images of *Slc32a1*, *Gad1*, and *Gad2* mRNA in LID mice. ISH signal intensities of *Slc32a1* (n=3), *Gad2* (n=4), and *Gad1* (n=4) mRNA compared between the Contra and Ipsi CPu, GPe, and SNr in LID and control mice.

(B) Expression levels of *Slc32a1*, *Gad1*, and *Gad2* mRNA were measured in the striatum of TD (n=5) and control (n=5) mice using quantitative reverse transcription polymerase chain reaction.

(C) AAV vector with ALFA-VGAT was injected into the right dorsal striatum of WT mice. Axon diameters and myelin thickness in the GPe and SNr were compared between the AAV injection (AAV Inj) and non-injection (AAV Un-inj) hemispheres (n=5).

(D) AAV vector with VGAT shRNA was injected into the right dorsal striatum of WT mice. Axon diameters and myelin thickness in the GPe and SNr were compared between the AAV Inj and AAV Un-inj hemispheres (n=5).

(E) A mixture of three AAVs fused with the tag proteins ALFA, 1D4, and HA (ALFA-VGAT, GAD67-1D4, and GAD65-HA) was injected into the right dorsal striatum of WT mice. Representative macroscopic images are of ALFA, HA, and 1D4 immunohistochemistry in the CPu, GPe, and SNr; representative SRM images are from the GPe.

(F) Representative SRM images of VGAT and PV immunohistochemistry in the AAV Inj and AAV Un-Inj hemispheres of WT mice with overexpression. The area of VGAT^+^ puncta of MSN terminals and the soma area of PV^+^ neurons in the GPe and SNr were compared between AAV Un-Inj and AAV Inj (n=4).

(G) Representative SRM images of PLP immunohistochemistry in WT mice with overexpression. The axon diameter and myelin thickness of cortical myelinated axons in the GPe and SNr were compared between AAV Un-Inj and AAV Inj (n=4).

*p<0.05, **p<0.01, ***p<0.001 (paired or Student’s t-test, p-values corrected by Bonferroni correction). Values are plotted as the mean ± SEM.

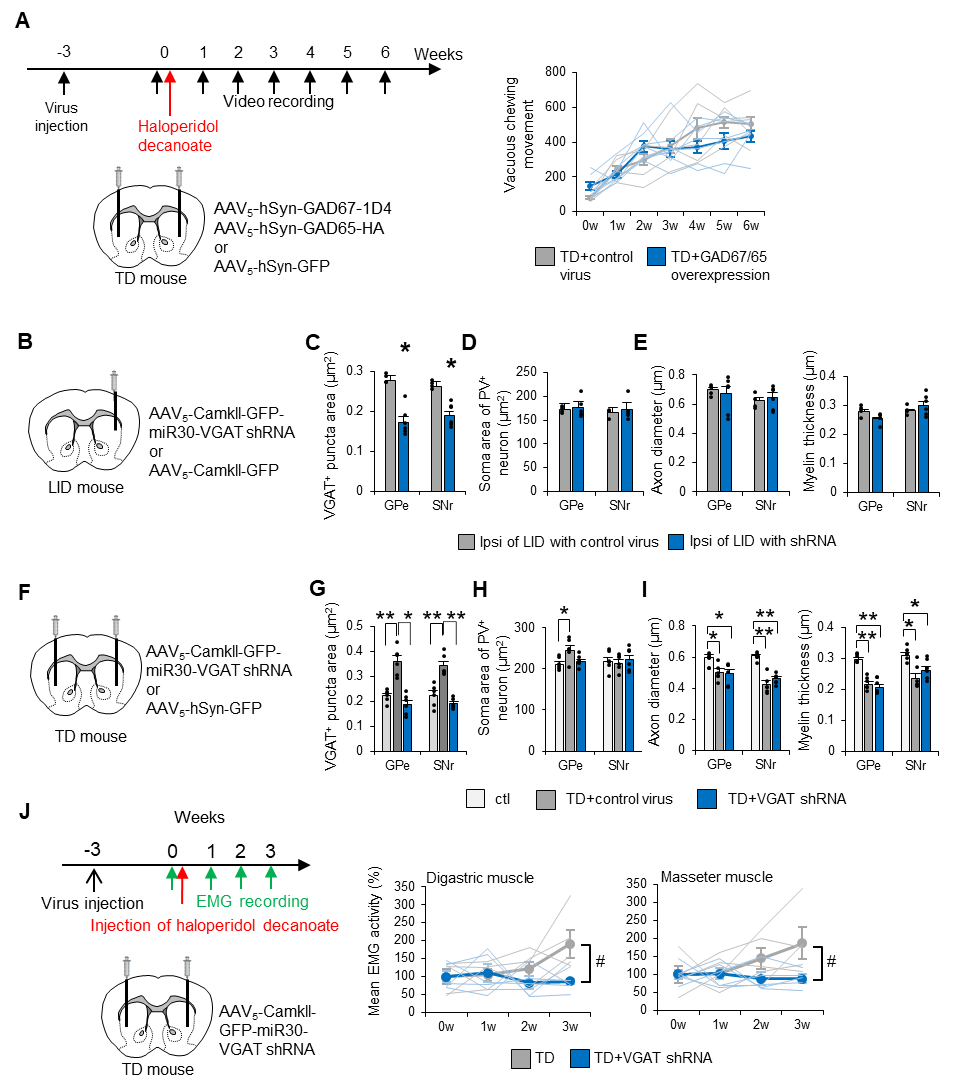

**Figure S5. VGAT inhibition suppresses VGAT^+^ MSN terminal enlargement but not GPe soma or cortical myelinated axon enlargement, related to Figure 4.**

(A) AAV vectors (a mixture of GAD67-1D4 and GAD65-HA, or GFP for the control) were injected into the bilateral dorsal striatum of WT mice 3 weeks before haloperidol decanoate administration. The number of VCMs was compared between TD mice with control AAV (n=6) and TD mice with GAD65/67 overexpression (n=6).

(B) AAV vectors (VGAT shRNA or GFP) were injected into the right dorsal striatum (ipsilateral to the 6-OHDA injection) of LID mice 2 weeks before the 6-OHDA injection.

(C–E) VGAT^+^ puncta area of MSN terminals (C), soma area of PV^+^ neurons (D), and axon diameter and myelin thickness of cortical myelinated axons (E) in the Ipsi hemisphere were compared between LID mice with control AAV (n=4) and those with VGAT-shRNA (n=5).

(F) AAV vectors (VGAT shRNA or GFP) were injected into the bilateral dorsal striatum of TD mice 3 weeks before haloperidol decanoate injection.

(G–I) VGAT^+^ puncta area of MSN terminals (G), soma area of PV^+^ neurons (H), and axon diameter and myelin thickness of the cortical myelinated axons (I) were compared among control mice (n=6), TD mice with VGAT shRNA (n=6), and TD mice with control AAV (n=6).

(J) AAV vectors (VGAT shRNA) were injected into the bilateral dorsal striatum of TD mice 3 weeks before haloperidol decanoate injection. EMG was recorded from digastric (jaw-opening) and masseter (jaw-closing) muscles for 3 weeks. Mean EMG activity from the digastric and masseter muscles was compared between TD mice with VGAT shRNA (n=5) and TD mice (n=5, same data as Figure S1F).

\# p<0.05 (two-way repeated ANOVA). *p<0.05, **p<0.01, ***p<0.001 (Student’s t-test, p-values corrected by Bonferroni correction). Values are plotted as the mean ± SEM.

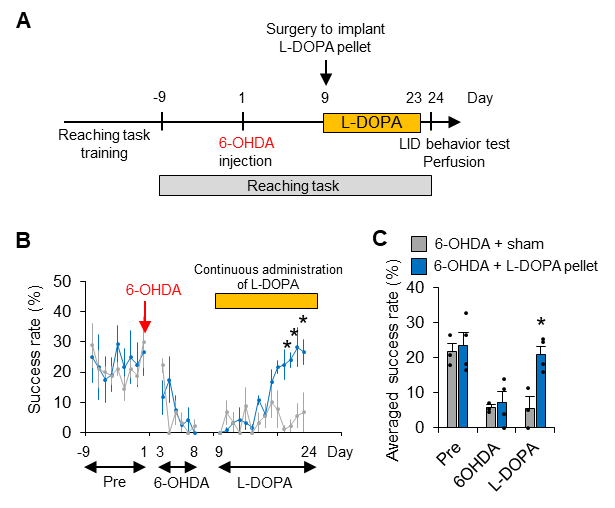

**Figure S6. Failure to produce dyskinesia is not the result of insufficient L-DOPA treatment, related to Figure 5.**

(A) Experimental time course for the model of continuous L-DOPA administration and reaching task. Mice were trained to perform the reaching task before the 6-OHDA injection, and the L-DOPA pellet was then implanted for continuous administration of L-DOPA. A sham operation was performed in control mice with 6-OHDA. LID behavior was measured after 2 weeks of L-DOPA administration.

(B, C) The success rate (B) and averaged success rate (C) in the reaching task were plotted for three periods: before injection of 6-OHDA (Pre), after injection of 6-OHDA (6OHDA), and after L-DOPA administration (L-DOPA).

*p<0.05 (Student’s t-test, p-values corrected by Bonferroni correction). Values are plotted as the mean ± SEM.

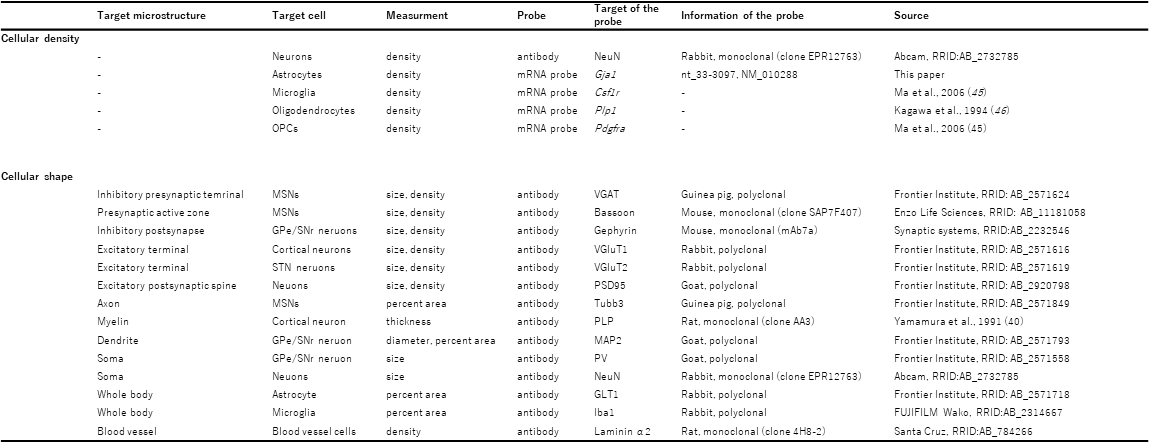

**Table S1. Comprehensive histological panel of brain anatomical investigations, related to Figure 1.**

Antibodies and mRNA probes for ISH, which were used to investigate contributions to brain volume changes, are listed.

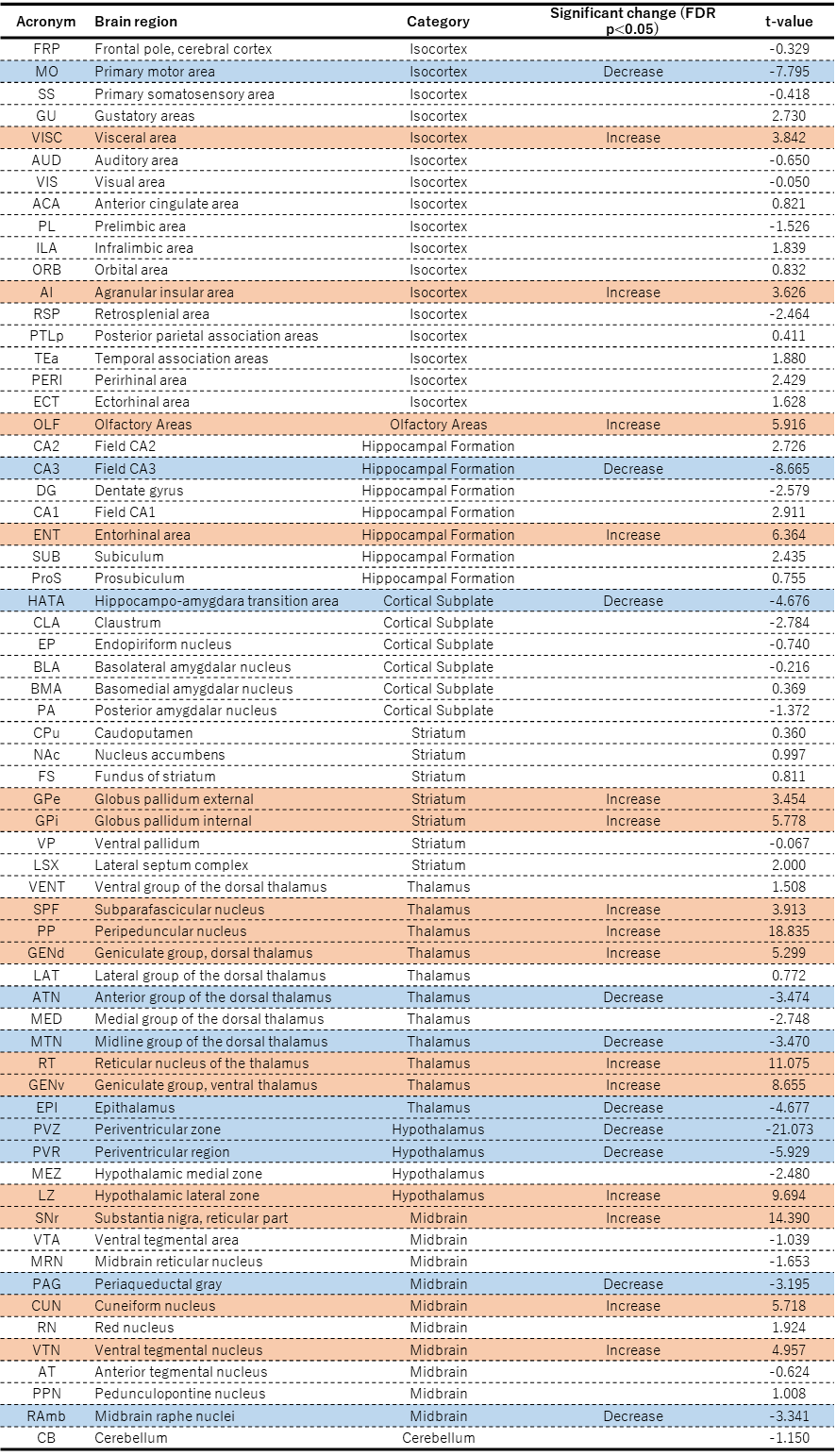

**Table S2. Summary of brain volume changes detected by ROI-based morphometry analysis of MRI in the LID model mice, related to Figure 1.**

Abbreviations, brain regions, and categories of brain regions were defined using the Allen Brain Atlas. There were significant volume changes (FDR-corrected p<0.05) in the ipsilateral hemisphere of LID model mice (n=8) compared with the findings in the contralateral hemisphere; t-values are shown. Red and blue indicate significant volume increases and decreases, respectively, in the ipsilateral hemisphere. FDR, false discovery rate.

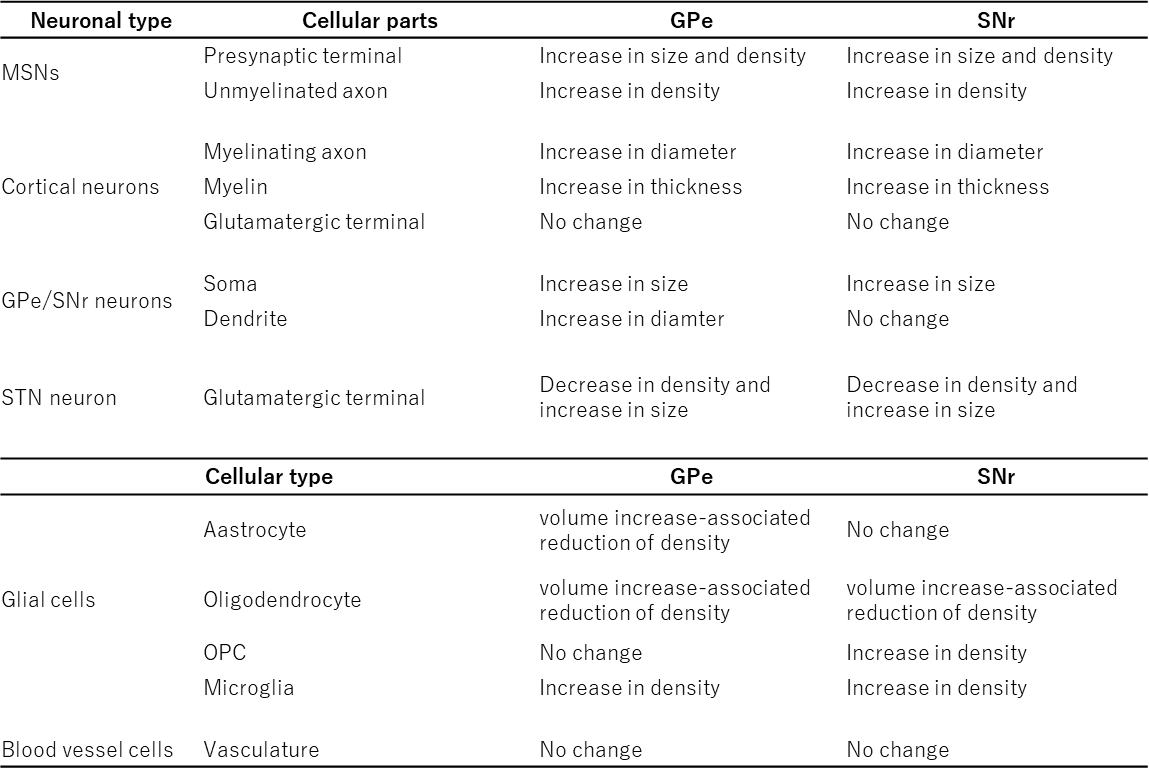

**Table S3. Summary of the contributions of structural changes in LID model mice, related to Figure 1.**

GPe, external segment of the globus pallidus; MSN, medium spiny neuron; OPC, oligodendrocyte precursor cell; SNr, substantia nigra pars compacta; STN, subthalamic nucleus.
